# Shapes and Genescapes: Mapping Multivariate Phenotype-Biological Process Associations for Craniofacial Shape

**DOI:** 10.1101/2020.11.12.378513

**Authors:** J. David Aponte, David C. Katz, Daniela M. Roth, Marta Vidal Garcia, Wei Liu, Fernando Andrade, Charles C. Roseman, Stephen A. Murray, James Cheverud, Daniel Graf, Ralph S. Marcucio, Benedikt Hallgrímsson

## Abstract

Realistic mappings of genes to morphology are inherently multivariate on both sides of the equation. The importance of coordinated gene effects on morphological phenotypes is clear from the intertwining of gene actions in signaling pathways, gene regulatory networks, and developmental processes underlying the development of shape and size. Yet, current approaches tend to focus on identifying and localizing the effects of individual genes and rarely leverage the information content of high dimensional phenotypes. Here, we explicitly model the joint effects of biologically coherent collections of genes on a multivariate trait—craniofacial shape — in a sample of *n* = 1,145 mice from the Diversity Outbred (DO) experimental line. We use biological process gene ontology (GO) annotations to select skeletal and facial development gene sets and solve for the axis of shape variation that maximally covaries with gene set marker variation. We use our process-centered, multivariate genotype-phenotype (MGP) approach to determine the overall contributions to craniofacial variation of genes involved in relevant processes and how variation in different processes corresponds to multivariate axes of shape variation. Further, we compare the directions of effect in phenotype space of mutations to the primary axis of shape variation associated with broader pathways within which they are thought to function. Finally, we leverage the relationship between mutational and pathway-level effects to predict phenotypic effects beyond craniofacial shape in specific mutants. We also introduce an online application which provides users the means to customize their own process-centered craniofacial shape analyses in the DO. The process-centered approach is generally applicable to any continuously varying phenotype and thus has wide-reaching implications for complex-trait genetics.

## Introduction

Variation in human craniofacial shape is moderately to highly heritable (∼30-70% (Cole et al., 2017; Tsagkrasoulis et al., 2017)), and resemblances among close relatives as well as twins underscore the strong relationship between shared genetics and shared phenotype(Johannsdottir et al., 2005; Nakata, 1985). Despite many studies in humans and in mice (Claes et al., 2018; Cole et al., 2016; Shaffer et al., 2016), however, we know very little about the genetic basis for variation in craniofacial shape. This is likely due to genetic complexity (Katz et al., 2019; Richtsmeier and Flaherty, 2013; Visscher, 2008; Wood et al., 2014; Wray et al., 2013). Like many aspects of morphological variation, craniofacial shape is extraordinarily polygenic. Genes with major mechanistic roles in facial development such as *Fgf8* often contribute little to standing phenotypic variation(Green et al., 2017) while genetic influences without obvious connections to craniofacial development emerge as significant contributors(Kenney-Hunt et al., 2008; Klingenberg and Leamy, 2001; Maga et al., 2015; Pallares et al., 2015, 2014). The effects of genetic variants on phenotype often depend on genetic background (Mackay and Moore, 2014; Percival et al., 2017) and many mutations have variably penetrant effects even when background is controlled (Hallgrimsson et al, 2009; Rendel, 1967). These issues likely arise because genetic influences act through multiple layers of interacting developmental processes to influence phenotypic traits, resulting in complex patterns of epistasis and variance heterogeneity(Hallgrimsson et al., 2018, 2014; Kawauchi et al., 2009; Wagner and Zhang, 2011). Solutions that go beyond studies of single gene effects are required to overcome these significant challenges in complex-trait genetics. Here, we implement an enhanced form of the more general candidate gene approach to evaluate the conjoint effects of multiple genes on a complex trait – craniofacial shape.

There are two basic approaches to mapping genetic effects on to phenotypic variation. A candidate gene approach measures genotypic values with known physiological and biochemical relationships to the phenotypes of interest (Cheverud and Routman, 1993). In contrast, a random marker or genome-wide approach seeks to associate any potential genetic variant with variation in the trait of interest. There are advantages and disadvantages to these two approaches. The candidate gene approach is blind to the unknown – phenotypic variation is often associated with genes not expected to be important. On the other hand, a candidate gene approach allows direct measurement of genotypic values and produces results that are interpretable in terms of trait physiology or development. A genome-wide or random marker approach can produce unexpected insight by revealing novel gene-phenotype associations. However, this comes at a great cost in power (Visscher et al., 2017). For highly polygenic traits, this approach often produces a “tip of the iceberg” effect in which studies reveal a small and often incoherent subset of the genes that actually determine variation in the trait of interest (Broman and Sen, 2009, p. 123-124).

Several strategies have been developed that partially overcome these tradeoffs. One solution is the use of polygenic risk scores. Polygenic risk scores assess the overall genetic influence on a trait without regard to the genome-wide significance of individual SNP effects (Dudbridge, 2013; Wray et al., 2007). Approaches such as meta-analyses of genome-wide association studies (GWAS) or studies based on extreme phenotypes (Morozova et al., 2015) have expanded gene lists for a variety of complex traits. However, lengthy lists of genes or overall genomic risk for specific phenotypes do not necessarily constitute tractable genetic explanations for phenotypic variation. When 1000s of genes are required to explain heritable variation in stature, for instance, it is not clear what such lists tell you beyond the obvious fact that stature is heritable and polygenic (Yang et al., 2010; Wood et al., 2014). This tension between hypothesis-driven and hypothesis-free approaches and their attendant tradeoffs between statistical power and interpretability is, arguably, a major issue within complex trait genetics. To resolve this conceptual conflict, approaches are needed that integrate quantitative genetics with biological insights regarding the cellular and developmental processes through which genes influence phenotypic variation.

Existing approaches to complex trait genetics also tend to treat phenotypic traits as singular and one-dimensional. Even for morphological variation, most studies reduce shape variation to linear distances, principal components, regression scores or measures of size which are then mapped as individual traits (Xiong *et al*. 2019; Shaffer et al, 2016; Cole et al, 2016).

This approach disregards the information content of multivariate phenotypic variation. While univariate traits only vary along one dimension, high dimensional traits such as craniofacial shape can vary in direction as well as magnitude within a multi-dimensional shape space. To identify the distinctive axes of gene effects on a multivariate trait, one must model such multiple multivariate relationships directly.

Building on Mitteroecker *et al.’s* (2016) multivariate genotype-phenotype (MGP) method, we extend the candidate-gene framework to evaluate the combined contributions of genes to variation in high-dimensional phenotypic traits such as craniofacial shape. Grouping genes by ontological information such as membership in pathways or other relevant biological hypotheses, our process-centered, multivariate approach brings traditional GWAS together with a simplified model of the hierarchical genotype-phenotype (GP) map. GP maps describe the relationship between genetic and phenotypic measurements(Lewontin, 1974). Understanding the genetic determinants of craniofacial variation, as with most complex traits, represents a many-to-many GP map problem (Fig 1). Both phenotypic and genotypic measurements have complex within-set covariance structures. On the genetic side, the covariance structure is represented by pathway/biochemical interactions, as well as chromosomal structure like linkage, chromatin, and 3D chromosomal organization. For shape-related phenotypes, the covariance matrix is structured by the chosen set of landmarks and their resulting coordinates. The functional relationship from genotype to phenotype is then described by a between-set covariance (Klingenberg and Leamy, 2001; Mitteroecker et al., 2016). To dissect these relationships, we use a regularized partial least squares (PLS) (Lorenzo et al., 2019) approach to estimate a low-dimensional mapping from the alleles in our sample to variation in adult mouse craniofacial shape. While PLS is well suited for analysis of covariation between two sets of measurements, regularization is essential for mitigating overfitting when there are many alleles simultaneously modelled. We focus on how allelic variation in processes relevant to craniofacial development maps to craniofacial shape variation. We ask the following five questions:

1. How much shape variation is communally accounted for by genes contributing to a process, e.g., chondrocyte differentiation?
2. How similar are the effects of different processes on shape? For instance, do cell proliferation genes affect face shape in a similar way to genes in the bone morphogenic protein pathway?
3. What is the overarching structure of process effects? Do process effects align with major axes of variation such as allometry or other principal directions in morphospace?
4. How similar are mutant model effects and process effects? For example, do chondrocyte mutant effects align with the effects of natural variants in chondrocyte differentiation genes?
5. Can one use the similarity of a mutational effect to MGP process effects predict unobserved phenotypes associated with that mutation?

**Fig 1.**
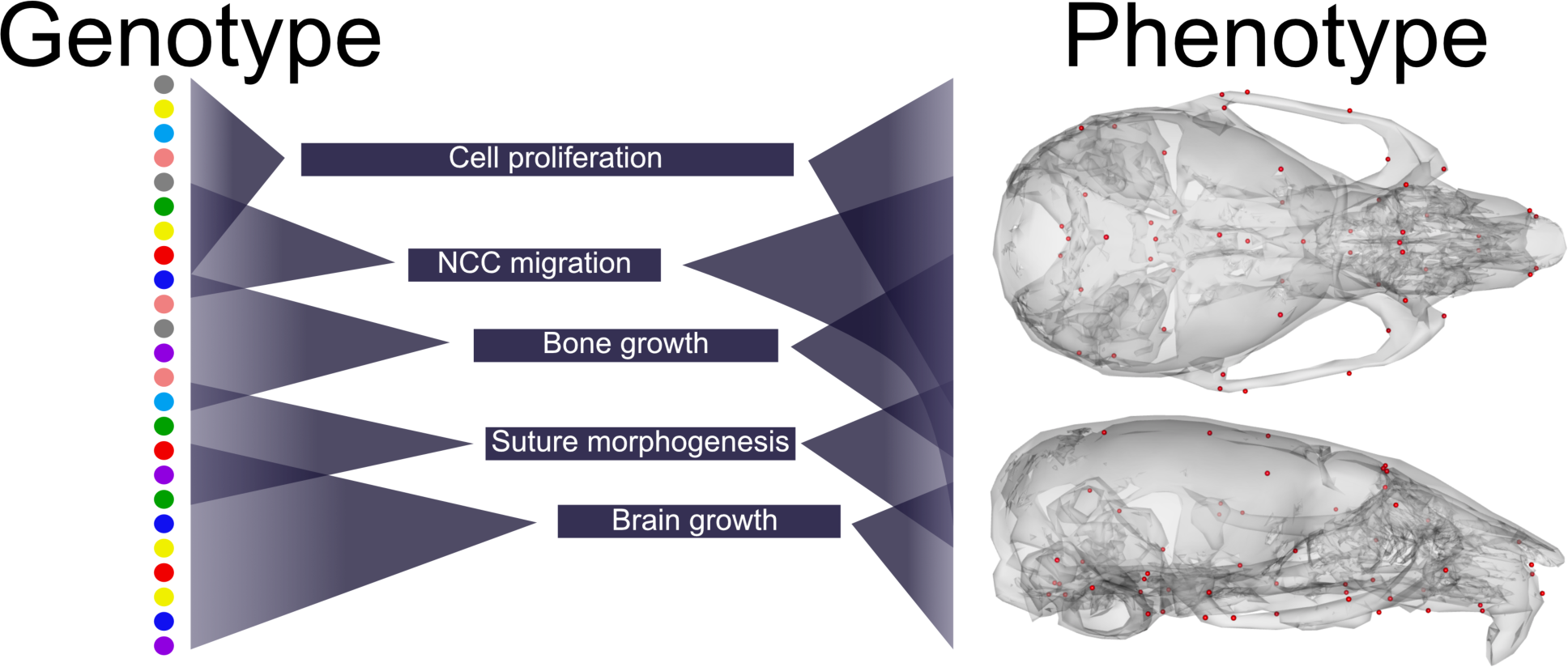
Schematic of the many-to-many relationship between genetic and phenotypic variation. From left to right: Allelic variation (colored dots) at individual genes is organized into developmental processes. Processes differ in start/end and duration during development. Genes are reused for different processes at different times. Processes are substantially pleiotropic in their effects contributing to global variation as well as local variation.

Together, these questions demonstrate the ability of the MGP approach to add meaningful understanding of the complex relationships between genotype and phenotype by quantifying higher level regularities between complex phenotypic and genomic data. We also demonstrate its potential as a resource for the study of mutational effects on complex traits such as craniofacial shape.

## Results

### Process Multivariate Genotype-Phenotype (MGP) Mapping

We demonstrate regularized-PLS MGP mapping with three examples. The first estimates the primary axis of skull shape covariation with genes involved in chondrocyte differentiation (Fig 2). Differentiation of chondrocytes is one of several key developmental processes involved in endochondral ossification. Endochondral bones form the majority of the cranial base through a cartilage model of bone formation (Percival and Richtsmeier, 2013). There are 38 genes annotated to chondrocyte differentiation in the Ensembl database (Yates et al., 2020). In the figure, genetic effects are shown as zero-centered bars that span the range of estimated allele effects across the 8 DO founders; individual founder allele effects—8 per marker—are color-coded within those bars (Fig 2A). Among chondrocyte differentiation genes, *Nov*, *Mapk14*, and *Bmpr1b* (*Alk6*) are most implicated in the major axis of pathway covariation with craniofacial shape. The phenotypic effects at each landmark—magnified 4x— primarily relate to antero-posterior positioning of the zygomatic arches and dorso-ventral jugal position (Fig 2B, 2C). The chondrocyte differentiation GP map explains 2.15% of the total variance in craniofacial shape. Compared to 1000 randomly generated marker sets of the same size (38), chondrocyte differentiation explains substantially more variation in phenotype than random markers (Supp fig 1A).

**Fig 2.**
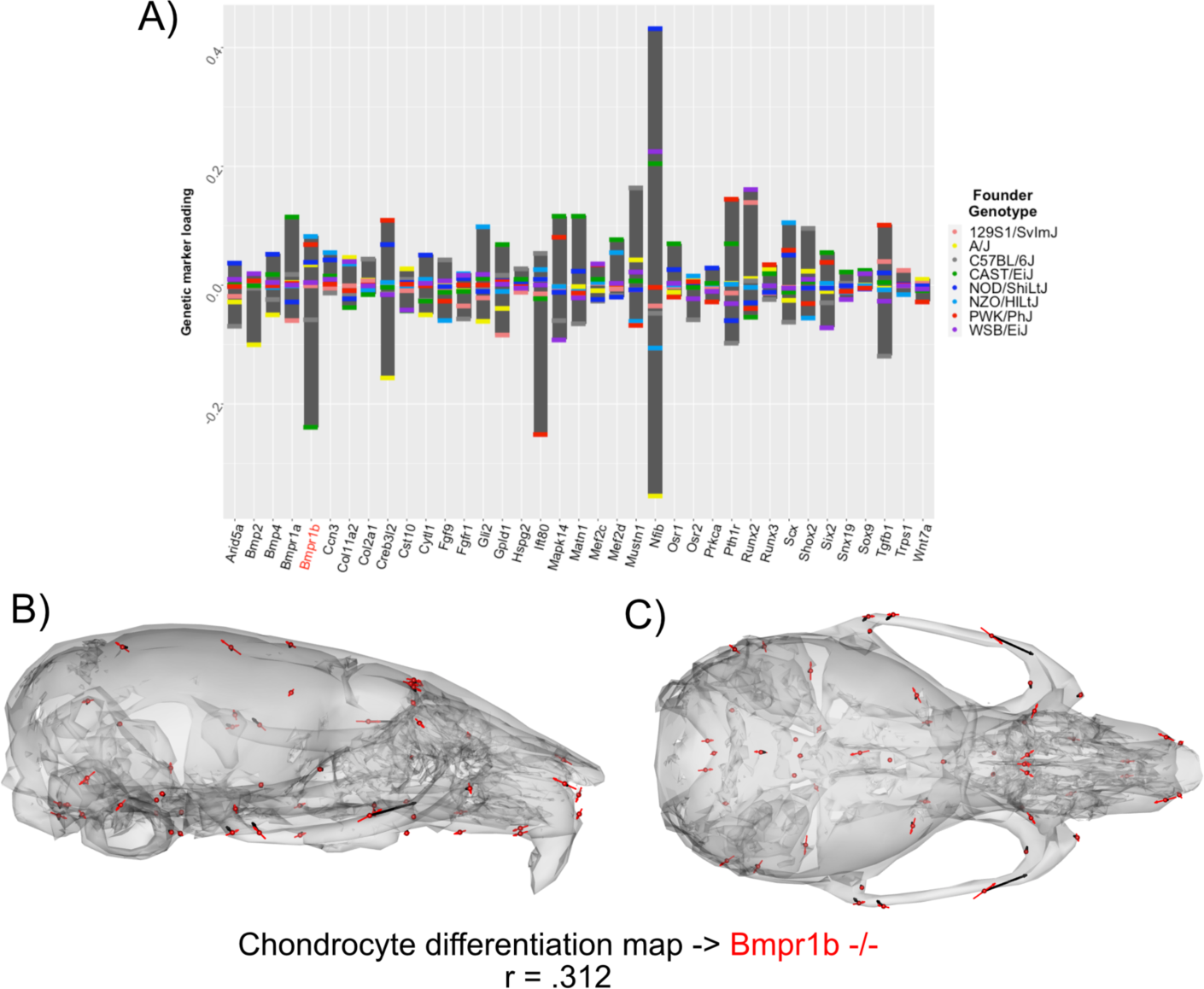
Process MGP for chondrocyte differentiation. A) PLS1 genetic loadings are shown for each gene in the model. Individual founder allele effect sizes are colored within each bar. B-C) Estimated chondrocyte differentiation MGP phenotype is shown with black vectors multiplied 4x. A *Bmpr1b* (*Alk6*) homozygous mutant is shown with red vectors for comparison. The vector correlation between chondrocyte differentiation MGP and *Bmpr1b* is shown below the phenotypic effects.

Figure 2B and 2C also compare the direction of the chondrocyte differentiation MGP axis to the axis of shape variation of a relevant mutant phenotype. We chose homozygous *Bmpr1b* mutants for this comparison for two reasons. The first is because Bmpr1b heterodimerization with other bone morphogenic protein pathway receptors is essential for chondrocyte differentiation and proliferation (Liu et al., 2005; Yoon et al., 2005). The second reason we chose *Bmpr1b* mutant comparisons is because the marker selected for *Bmp1rb* in the genomic analysis is contains one of the strongest allelic effects associated with the morphological effect. The overall phenotypic directions of *Bmpr1b* mutant variation and chondrocyte differentiation variation are moderately correlated at r = 0.312, but the direction at landmarks with large effects in mutant and MGP are clearly coincident. Over the landmarks we measured, the chondrocyte differentiation effect is less global than the *Bmpr1b* effect, likely due to the difference in severity of the mutant phenotype.

The similarity of the chondrocyte differentiation effect with the *Bmpr1b* mutant and the high loading *Bmpr1b* allele in the DO genome suggests that *Bmpr1b* mutants may produce chondrocyte differentiation defects in the developing neurocranium. We quantified cell size and distribution in the intersphenoid synchondroses (ISS) of several mutant and control *Bmpr1b* mice. Homozygotes show overall larger cell sizes as well as a differing distribution of cell sizes throughout the width of the ISS (Fig 3A-C; χ^2^ = 21.23, df = 3, p < .0001). The presence of larger cell sizes in the homozygote *Bmpr1b* mutants suggests that the synchondroses possess more hypertrophic chondrocytes. Additionally, *Bmpr1b* homozygous mutant mice show premature fusion of the coronal suture (Fig 3D).

**Fig 3.**
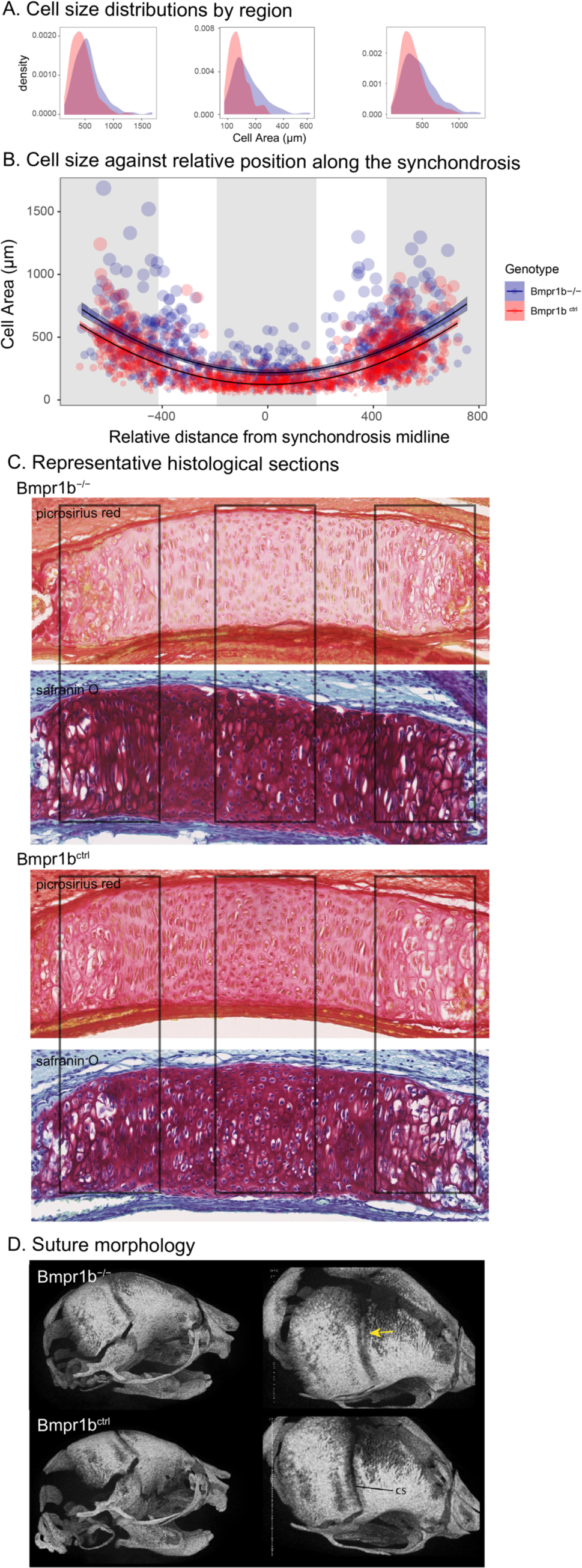
Chondrocyte defects in *Bmpr1b* mutants. A-B) Quantification of cell size in the sections of the intersphenoid synchonrosis shows an increase in relative cell size as well as a change in the distribution of cell sizes throughout the width of the synchondrosis. C) Sections of intersphenoid synchondroses. D) Premature fusion of the coronal suture is visible in *Bmpr1b* homozygous mutants.

The second example quantifies cranial shape covariation with the 81 genes annotated to “determination of left/right symmetry”. The phenotype associated with left/right symmetry alleles is predominately related to a larger neurocranium volume relative to the outgrowth of the face (Fig 4B, 4C). We also visualized the asymmetry in the phenotypic response, which shows subtle asymmetry, particularly in the position of the anterior zygomatic landmark (Fig 4D).

**Fig 4.**
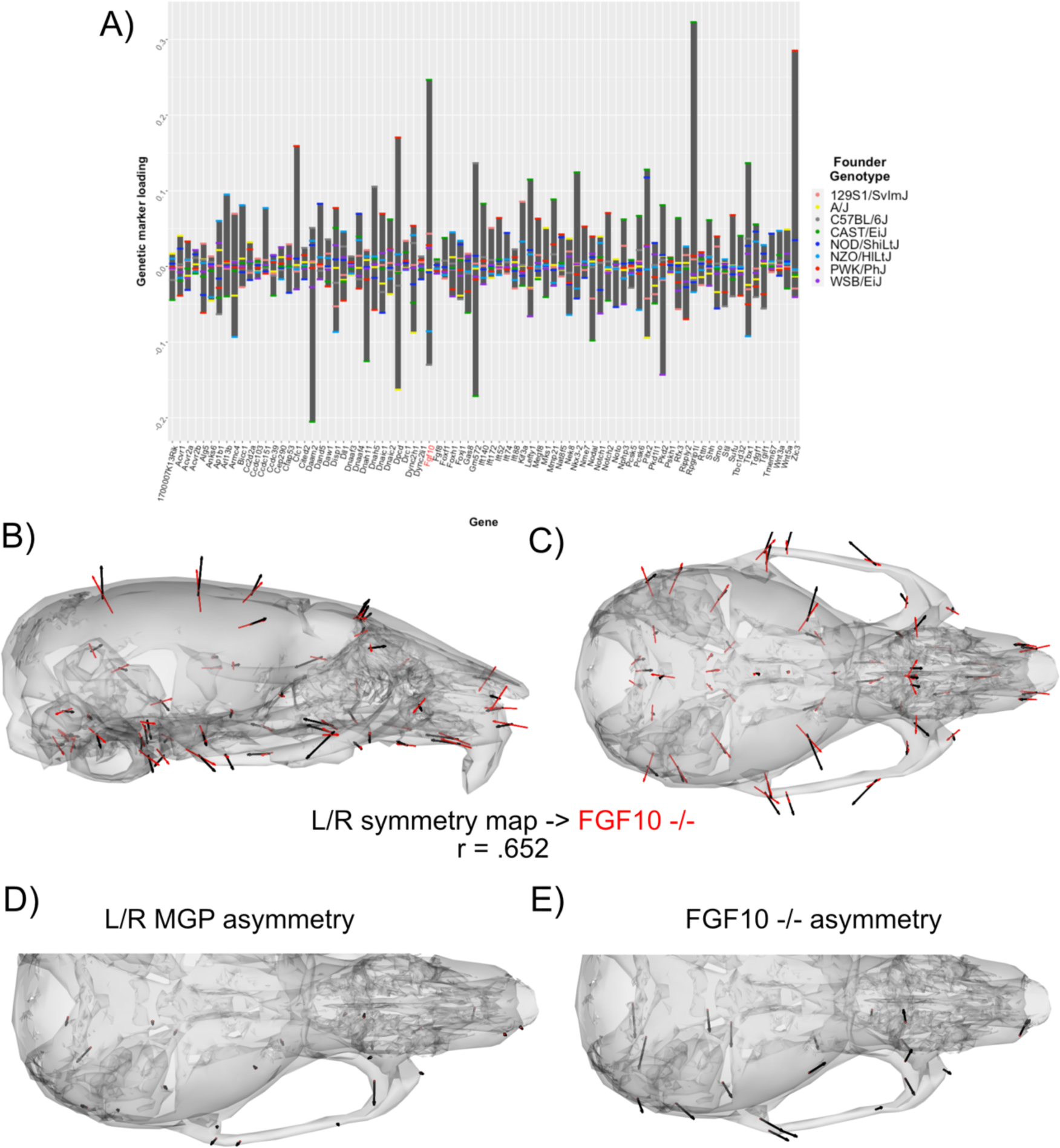
Process MGP for determination of left/right symmetry. A) PLS1 genetic loadings are shown for each gene in the model. Individual founder allele effect sizes are colored within each bar. B-C) Estimated left/right symmetry MGP phenotype is shown with black vectors multiplied 4x. An *Fgf10* homozygous mutant is shown with red vectors for comparison. The vector correlation between left/right symmetry MGP and the *Fgf10* mutant is shown below the phenotypic effects. D-E) Visualizations of asymmetry in the L/R MGP response and the *Fgf10* homozygous mutant. Asymmetry vectors are magnified 4x.

Left/right symmetry loci explain 2.2% of the total variance in craniofacial shape, which exceeds the variance explained by 1000 randomly selected marker sets of the same size (Supp fig 1B).

There are several high loading alleles that contribute to the left/right symmetry phenotype. In particular, an *Fgf10* allele inherited from the Castaneus founder background was among the most important (Fig 4A). FGF10 is a key ligand in early development, directing proliferation as well as differentiation for many craniofacial components, including the palate, teeth, and bones (Hilliard et al., 2005; Prochazkova et al., 2018; Watson and Francavilla, 2018). We compared the estimated left/right symmetry MGP effect with the direction of an *Fgf10* homozygous mutant because of the relative importance of the allelic effect. The vector correlation between the *Fgf10* mutant and the estimated left/right symmetry effect is 0.63.

The high-loading *Fgf10* allele for left/right symmetry along with the similar genomic and mutant phenotypes suggests that *Fgf10* mutants could show directional asymmetry in the cranium. To test this, we measured a sample of 8 *Fgf10* adult mutant crania for object symmetry and detected significant directional asymmetry (Fig 4E; F = 4.91, df = 52, p < .0001).

The final example estimates the shape covariation attributed to the 73 genes annotated to “palate development.” Formation and fusion of the palatal shelves are crucial for proper orofacial development and heavily influences overall facial shape (Greene and Pisano, 2010). Several genes contribute strongly to the palate development MGP effect including *Ephb2, Gli3,* and *Lrp6*. The estimated phenotype shows corresponding variation in palate length as well as strong effects in the majority of the cranial base landmarks (Fig 5B, 5C). Palate development MGP loci explain 2.4% of the total variance in cranial shape, which is greater than variance explained by 1000 randomly permuted marker sets of the same size (Supp fig 1C). We compared the palate development phenotype to a heterozygous *Ankrd11,* neural-crest specific knockout mouse. The *Ankrd11* locus is associated with KBG syndrome in humans, which presents with generally delayed bone mineralization as well craniofacial characteristics including palate abnormalities (Low et al., 2016). While the vector correlation between the palate development MGP effect and the *Ankrd11* mutant over the complete set of cranial landmarks is moderate at r = .284, the vector correlation for palate landmarks is substantially higher at r = .536.

**Fig 5.**
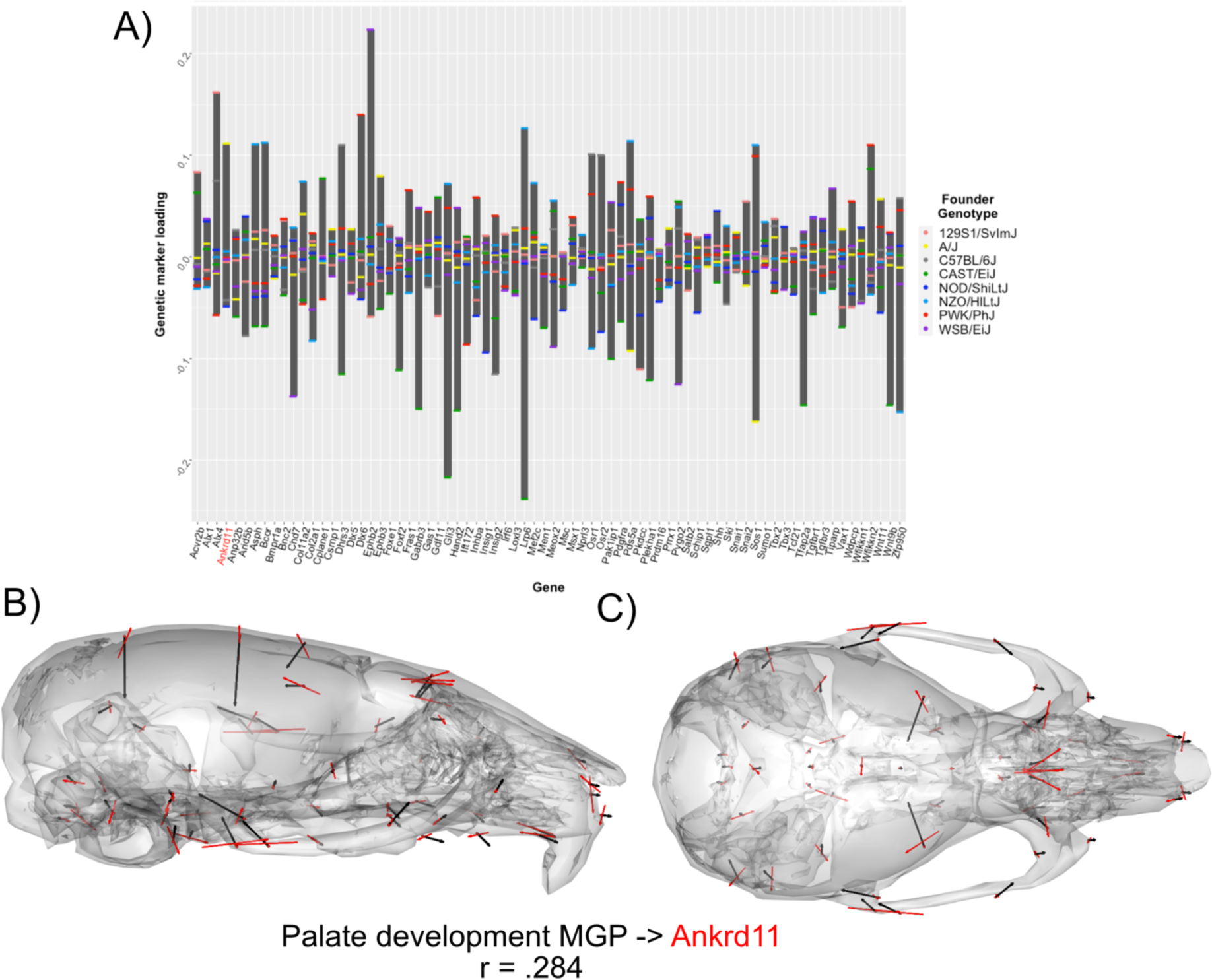
Process MGP for palate development. A) PLS1 genetic loadings are shown for each gene in the model. Individual founder allele effect sizes are colored within each bar. B-C) Estimated palate development MGP phenotype is shown with black vectors multiplied 4x. An *Ankrd11* mutant is shown with red vectors for comparison. The vector correlation between palate development MGP and the *Ankrd11* mutant is shown below the phenotypic effects.

In each case above, we have shown how association of gene sets and phenotypic variation can produce highly informative results that can guide future hypothesis testing. For a given biological process, we identified genes that load strongly on the primary axis of MGP covariation for which mutant samples were available to us, as well. Future investigations could also use this information about genes with high loadings to generate new mutants for analysis. For each example, we focus only on the first PLS axis, so other alleles for other genes may contribute to novel phenotypic directions in lower PLS axes. In the next sections we will examine how MGP phenotypes relate to each other, as well as the phenotypic directions of many mutant mouse models.

### Pairwise comparison of craniofacial development processes

We chose 15 processes integral to craniofacial development and compared the pairwise similarity of effect on craniofacial shape using a heatmap based on clustering of the correlation matrix (R core team, 2017). Processes with similar effects on craniofacial shape will be highly correlated, while processes that affect distinct aspects of craniofacial variation will be uncorrelated to each other. The clustering algorithm resulted in two main blocks of strongly correlated effects (Fig 6A). The largest block of highly correlated phenotypic effects includes neural crest cell migration, epithelial to mesenchymal transition, forebrain development, as well as some of the most general developmental processes like cell proliferation, bone development, apoptosis, A/P pattern specification, and FGFR signaling. In addition, there is a general BMP block, with Bmp signaling, dorsoventral pattern formation, endochondral ossification, and positive regulation of skeletal muscle tissue growth. Interestingly, phenotypic variation associated with cranial suture morphogenesis, neural tube patterning, and intramembranous ossification is largely uncorrelated with the other craniofacial developmental processes included here.

**Fig 6.**
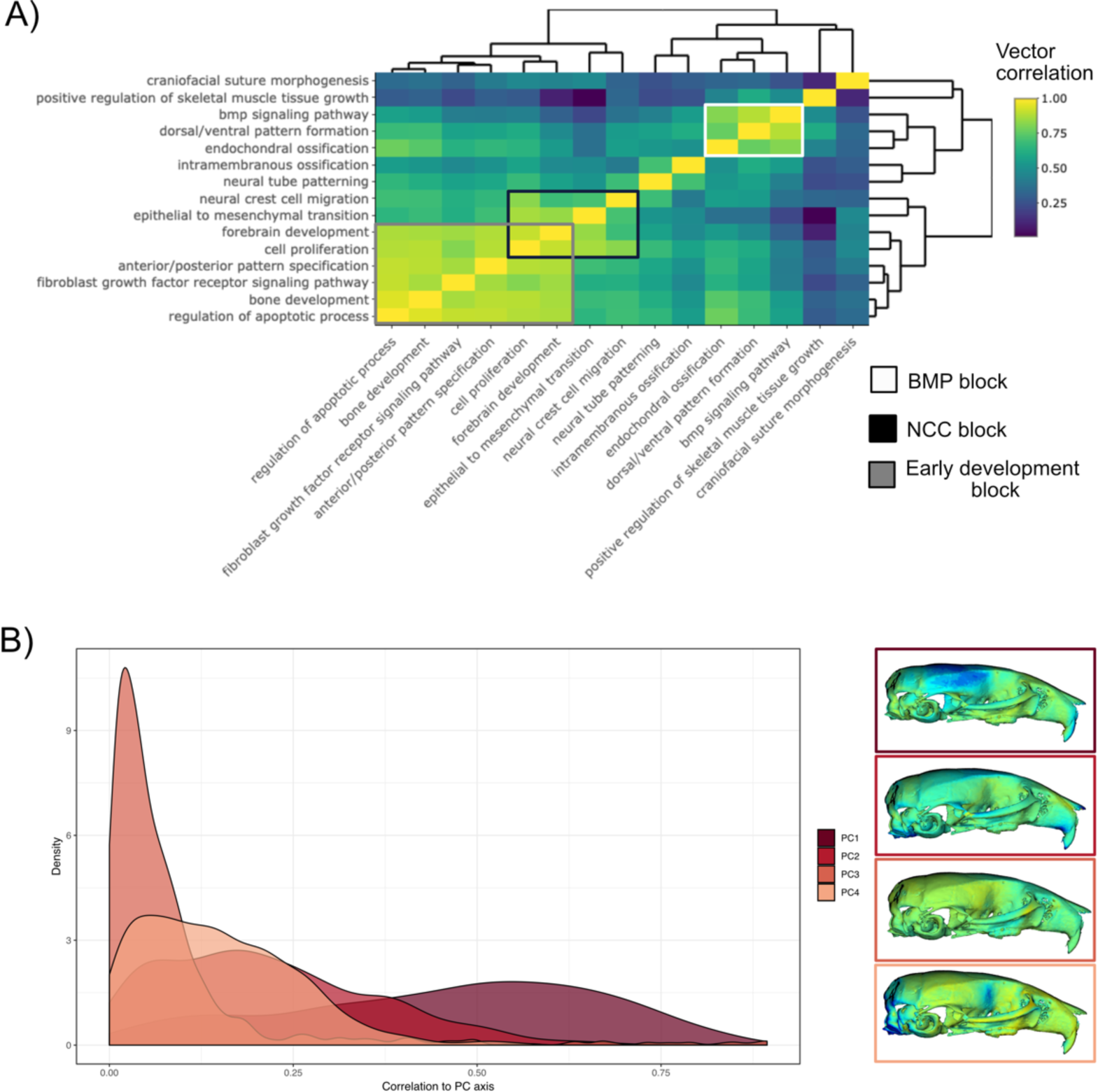
Pairwise MGP vector correlations and random process correlations to principal components. A) Pairwise correlations of phenotypic effects for 15 process MGP analyses. Scale on the right denotes color correspondences to vector correlation, where yellows are high correlations, greens are moderate, and blues are low. B) The densities of vector correlations between 1000 randomly chosen process MGP effects and PCs 1-4.

To assess the stability of the clustering result, we estimated the vector correlation between the cluster distances—also known as the cophenetic distance— and the original correlation matrix. A high vector correlation suggests reliable clustering, whereas a low correlation suggests a random clustering result. The correlation between the cophenetic distance matrix and the correlation matrix is 0.648 (t = 8.64, df = 103, p = 7.6^-14^), suggesting a moderate, though significant structure in the similarity of effects amongst this set of MGP processes.

### Comparison of processes to principal component directions

Almost a third of the 15 pairwise process comparisons showed a vector correlation > 0.5, suggesting that many processes may feed into a limited set of directions in morphospace. To assess the extent to which different processes affect the same aspects of facial shape we randomly chose 1,000 process annotations, fit individual regularized PLS models to each set of markers for a given annotation term, and then compared the direction of phenotypic effects for each model to principal components 1-4 of the DO shape data (Fig 6B). Doing so with principal components allows us to highlight similarities in directions of process effects. Process effects range from completely uncorrelated to PC1 to highly correlated (0.0 - 0.8). However, the central tendency of randomly selected process effects is one of moderate-to-high correlation with PC1. Moderate correlations (0.55 - 0.6) with PC1 are more common than uncorrelated effects.

Supplemental table 1 contains the 10 most highly correlated processes with PC1 as well as the corresponding correlations to PC 2-4. The most highly correlated process to PC1 is “zinc ion binding”, which is 0.86 correlated to the PC1 direction. The remaining 9 processes most highly correlated to PC1 includes “sensory perception of sound”, “Calcium ion transport”, “Protein homooligomerization”, “Dendrite morphogenesis”, “Neuropeptide signaling pathway”, “Focal adhesion”, “Chromosome segregation”, “Sarcomere organization”, and “Integral component of endoplasmic reticulum membrane”.

Process correlations with PCs 2-4 are generally less strong. The maximum correlated process with PCs 2-4 was “V(D)J recombination”, “n-terminal protein myristoylation”, and “branching morphogenesis of an epithelial tube” with vector correlations of 0.72, 0.89, and 0.56, respectively. Processes with high vector correlations for a given PC tend to be uncorrelated with other PCs (Supp table 1), although some processes load moderately high across several PCs. For example, “negative regulation of I-kappaB kinase/NF-kappaB signaling” shows vector correlations between 0.25 - 0.54 for the first four PCs (Supp table 2).

### Process effects in the mutant morphospace

To assess the extent to which craniofacial shape variation associated with developmental processes aligns with variation from mutants of major effect, we projected 7 process effects onto the first two principal components (PCs) of a dataset containing the DO sample, and samples from 30 mutant genotypes (Fig 7A). Each black label represents the mean shape score of the listed mutant genotype. The shaded ellipse with an orange border displays the 95% confidence ellipse of PCs 1 and 2 of DO cranial shape variation. The DO mean shape is contrasted by the mutant variation along PC1. The first PC describes vault size relative to the length of the face.

**Fig 7.**
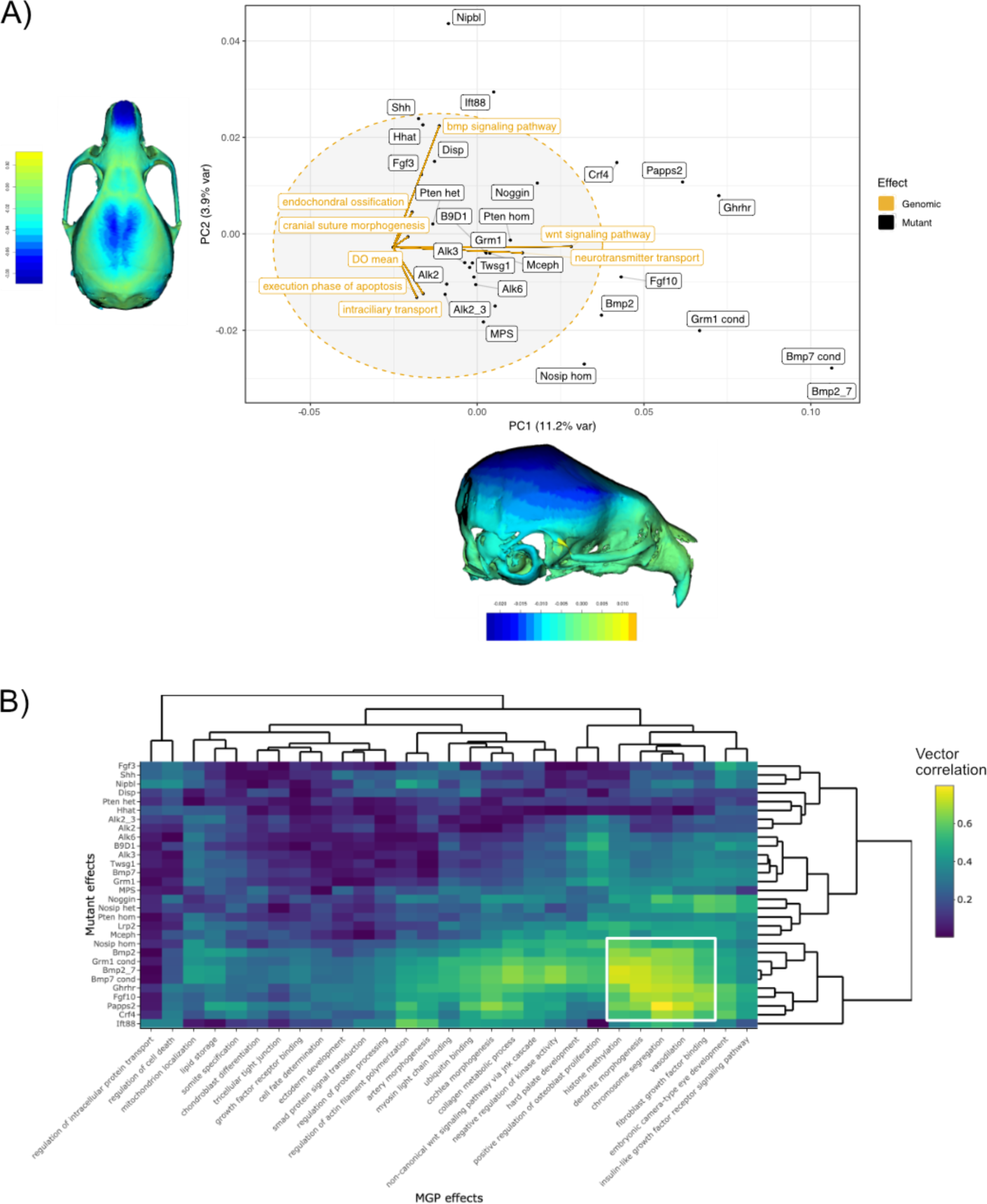
Comparisons of MGP and mouse mutant directions. A) Seven MGP phenotypes projected onto a PCA of the DO and a sample of 30 mutant mouse genotypes. Mutant means are labeled in black. The directions of MGP effects are shown with orange vectors from the DO mean to the associated process MGP. The range of DO variation on PCs 1 and 2 is shown with the shaded ellipse with an orange border. B) A heatmap of vector correlations between 30 mutant effects and 30 process MGP effects.

The phenotype shown along the x-axis of figure 7A depicts the maximum positive PC1 shape, while the heatmap drawn on the crania represents the local deformations towards the minimum PC1 shape. The positive direction of PC2 describes coordinated variation that includes a relatively wider vault, narrower zygomatic, and shorter premaxilla (Fig 7A, y-axis margin).

Process effects—highlighted with orange vectors originating at the DO mean shape — are necessarily of smaller magnitude than the total variation in the DO sample. Therefore, to better compare the direction of process effects the vector magnitudes were magnified 4x. Several process effects align in distinct directions of mutant effects, such as bmp signaling pathway and endochondral ossification in the direction of *Shh*, *Nipbl*, and *Ift88* mutants. Neurotransmitter transport and Wnt signaling pathway is similar in direction to *Mceph* and *B9d1* mutant effects. Execution phase of apoptosis and intracellular transport both show similar effects to a cluster of Bmp mutants.

Finally, we show the similarity of 30 process MGP effects to 30 mouse mutant models in figure 7B. The heatmap shows the correlation in direction with yellow/green denoting higher correlation and teal/blue denoting lower correlation. The bottom right of the heatmap (highlighted by a white border) shows a block of mutants for which there are strong process correlations. These are among the most extreme phenotypes along PC1 (Fig 5A) and include mutants for *Nosip, Bmp2, Grm1, Bmp2; Bmp7* transheterozygote*, Bmp7, Ghrhr, Fgf10, and Papps2*. The processes most strongly correlated to these mutants are histone methylation, dendrite morphogenesis, chromosome segmentation, vasodilation, and fibroblast growth factor binding.

There are a set of mutant phenotypes that have generally low correlations to the set of processes chosen. These mutants include *Fgf3, Shh, Nipbl, Disp, Pten, Hhat, and Alk2; Alk3* transheterozygote. Interestingly, this group of mutants vary more along PC2 than PC1 (Fig 7A). Notably, regulation of intracellular protein transport and regulation of cell death are strongly uncorrelated with the majority of mutant directions.

### Real-time process GP mapping

Finally, we provide an online tool to visualize process effects and make comparisons to mutant effects in real time. This application is found at **genopheno.ucalgary.ca/MGP** and can be used for analyses similar to those described in this paper. When the user selects gene ontology terms, the program searches for genotype markers adjacent to each gene listed and uses the selected markers to fit a regularized PLS model. The result is an estimate of the many-to-many relationship between the selected markers and cranial shape variation. The visual outputs include barplots depicting the relative allele effect sizes for each gene in the process and a 3D plot of the corresponding axis of shape variation. Users can compare the effects of different processes and also compare process effects to mutant effects from a provided database of 30 mutant genotypes.

To illustrate how to use this application, we have provided the graphical user interface used to select the parameters (Fig 8). As an example, in the “Process text” entry field, supply a starting term; we chose “brain.” The GO database is then filtered, returning a user-selectable subset of biological process ontology annotation terms in the “Process filter” field. We chose “forebrain morphogenesis,” which has 11 associated genes. We chose to magnify the process phenotype vectors 4x and compare the effect to a heterozygous *Ift88* mutant. *Ift88* is a core component of the primary cilia, which are responsible for promoting developmental signals involved in many facets of facial development (Tian et al., 2017). Further, the plots that are generated are interactive. For example, marker loadings can be highlighted and subset by genes of interest (Plotly, 2015). There is further information about using this online tool in the “About this app” tab.

**Fig 8.**
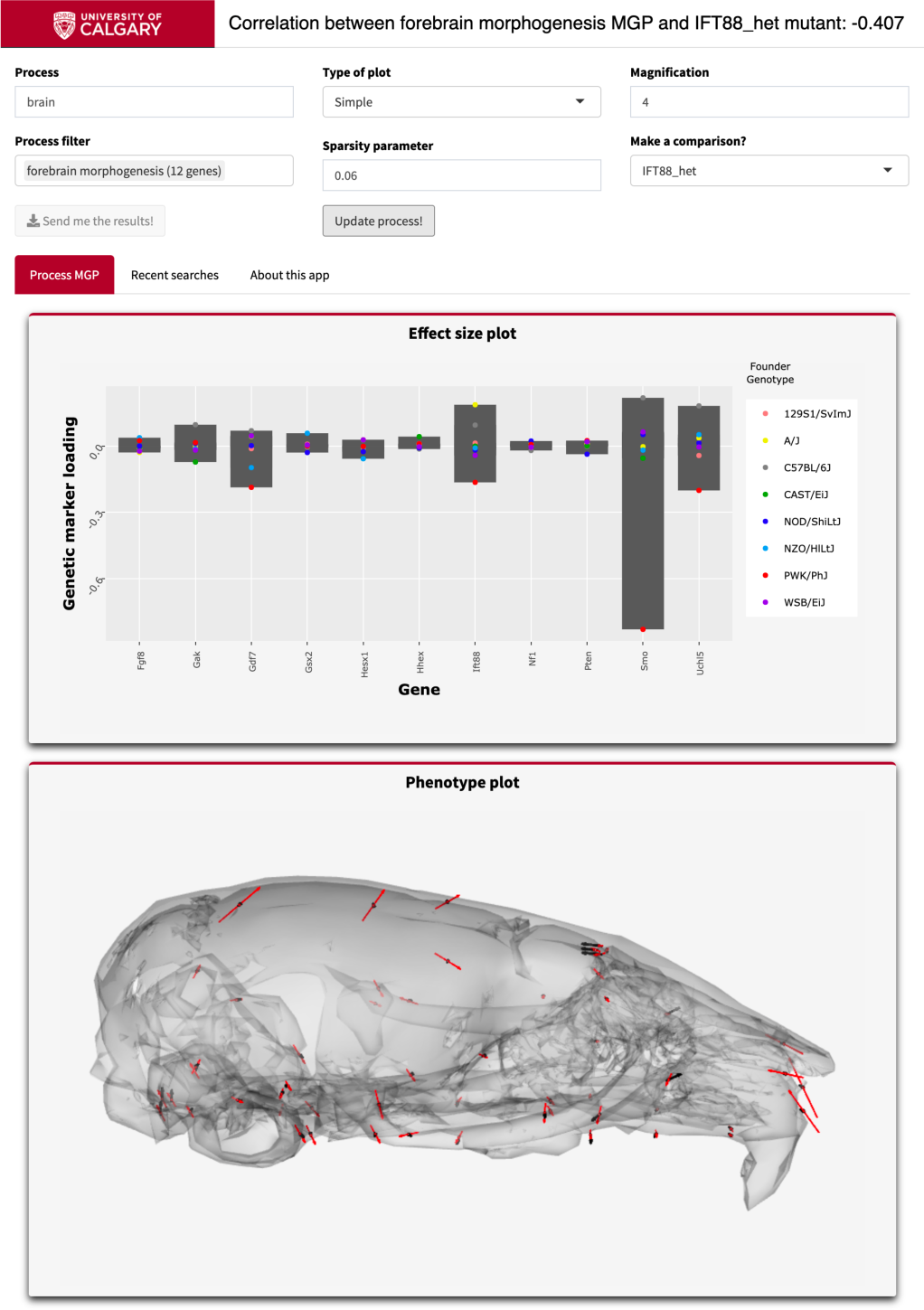
Example screenshot of web version of process analysis. Analyses include a barplot of the relative effect sizes of each selected marker and the associated phenotype shown with black vectors at each landmark. If a mutant comparison is selected, the vector correlation is provided and the mutant phenotype is shown with red vectors. Selecting “send me the results” generates an HTML report with an interactive 3D model.

## Discussion and Conclusion

A key goal in genomics is to create tractable genetic explanations for phenotypic variation. In this study, we used a regularized PLS approach to model the joint effects of genomic markers on multivariate craniofacial shape. This innovative approach allows us to address the joint contributions of multiple genes that share ontological characteristic such as pathway membership on craniofacial shape as a multivariate trait. Specifically, we chose markers adjacent to genes annotated under a developmental process of interest. We showed three process MGP analyses in depth, each with distinct phenotypic effects. The chondrocyte differentiation MGP effect mainly showed effects on the shape of the zygomatic and jugal bones with *Ccn3/Nov* as the most highly loaded corresponding marker effect. The left/right symmetry MGP phenotype was primarily a smaller cranial vault volume with a longer facial outgrowth, broadly similar to the primary axis of shape change during mouse growth and development (Gonzalez et al., 2013). The most highly loaded markers for the left/right symmetry effect were related to *Fgf10* and *Rpgrip1l.* We then compared process MGP phenotypic effects to each other, to mutant phenotypes, and the first four principal components of the diversity outbred sample. Each of these comparisons highlighted the integrated structure of phenotypic variation in mouse craniofacial shape. We found that while there are processes with distinct and localized effects, genetic effects generally converge on a limited set of directions in phenotype space. Further, these process effects often correspond with the directions of major mutations known to affect these same processes.

Many recent studies have addressed the genetics of craniofacial shape in humans and mice (reviewed in: Roosenboom et al., 2016; Weinberg et al., 2018). While these studies are yielding a growing list of genes, suggesting that facial shape is highly polygenic, they have left the vast majority of heritable variation unexplained. Existing studies have either used univariate measures of facial shape such as linear measurements or univariate summaries of multivariate shape (eg. Procrustes distances or PC scores). In addition, most genomic studies of craniofacial shape quantify the effects of each genomic marker independently, with notable exceptions focusing on epistatic effects (eg. Varón-González et al., 2019). Our approach shares common features with some predecessor GP mapping strategies in which candidate genes/SNPs are selected a priori because of common involvement in a pathway (or other mechanistic cluster) (Claes et al., 2014; Liu et al., 2012; Wang et al., 2010, 2007). In particular, Wang and colleagues selected SNPs based on proximity to genes of interest and effect size to jointly model the pathway-level effects on Parkinson disease data. Their approach is similar to gene-set enrichment analysis, weighing over-representation of statistical effects related to case-control group membership. In contrast, our approach focuses on estimating a multivariate continuous set of craniofacial responses. Importantly, our approach jointly identifies genotype-phenotype axes that maximally covary. This differs significantly from approaches that determine phenotypes for analysis *a priori* or based on a pre-determined method of data reduction such as PCA. Our approach also differs from methods that associate single locus effects with a multivariate phenotype (Claes et al., 2018).

A key finding of our application of the MGP method to craniofacial shape is that multivariate phenotypic variation aligns nonrandomly to genetic markers associated with pathways or developmental processes. For most process MGP maps, multiple markers for gene sets with known developmental relationships covary in their relationship with craniofacial shapes (Supp fig 2). These covarying effects represent joint genetic effects of multiple contributors to phenotypic variance. While these patterns of multivariate genotype-phenotype covariation may include genetic variants that do not actually affect the phenotype, many others will be contributors that we lack statistical power to detect under a typical univariate approach (Pitchers et al., 2019; Varón-González et al., 2019). Here, the overall pattern of genotype-phenotype covariance is the level of genetic explanation for phenotypic variation. When such patterns involve genes that are ontologically linked in meaningful ways, they provide a level of insight into the developmental-genetics of phenotypic variation that is beyond reach for most genome-wide association studies for complex traits.

Another valuable asset that arises from the MGP approach is the ability to generate testable hypotheses or predictions from multivariate genotype-phenotype observations. The chondrocyte differentiation analysis suggested differentiation defects in the *Bmpr1b* mutant. Subsequent histological analysis of *Bmpr1b* mutants showed premature suture fusion as well as atypical distribution of hypertrophic chondrocytes in the intersphenoid synchondrosis. Similarly, the MGP analysis of left/right symmetry genes suggested that *Fgf10* alleles can contribute to directional asymmetry. A follow up morphometric analysis of symmetry showed that *Fgf10* mutants do display significant craniofacial asymmetry (Fig 4E). MGP can also be used to test existing hypotheses about genotype-phenotype relationships. The relative importance of the *Ankrd11* locus in the palate development analysis and the similarity between the genomic and mutant phenotype further validates the role of Ankrd11 in palate development. These examples illustrate the additional insights that an MGP analysis of a mutational effect can provide. Given that such comparisons can be run quickly, this creates a tool with tremendous potential for hypothesis generation and initial screening for hypotheses about process-level effects.

In aggregate, our results show substantial covariance in the directions of phenotypic effects among different developmental processes (Figure 6A). The largest of these captures processes general to development such as cell proliferation or pattern specification. The second captures processes more specific to craniofacial development such as cranial suture/ossification and neural tube patterning. While processes are structured in their effects, our data suggest that many processes likely “add up” to produce variation. Thus, of 1000 randomly selected processes, 25.7% had a PC1 correlation higher than 0.6, supporting not only a highly polygenic model of facial variation, but one in which hundreds or even thousands of developmental processes that contribute to craniofacial variation. Importantly, this result shows how many processes and pathways converge to produce central axes of variation in craniofacial shape.

The explicit modeling of multivariate relationships between phenotypes and genotypes also allows a focus on pleiotropy. Developmental studies in mice demonstrate widespread craniofacial morphological effects from localized developmental perturbations (Martínez-Abadías et al., 2012; Stelzer et al., 2007; Young et al., 2010) Perturbations to specific processes in development generally produce effects on multiple aspects of phenotype due to knock-on effects at later stages or to interactions at the level of tissues or anatomical structures (Hallgrimsson et al, 2007). A change in cartilage growth in basicranial synchondroses produces a global change in craniofacial form, for example (Parsons et al, 2015). Remarkably, enhancers with highly specific temporospatial effects on gene expression also produce global rather than localized changes in craniofacial shape (Attanasio et al., 2013). Given that pleiotropy is likely ubiquitous (Hill and Zhang, 2012; Wagner et al, 2008), explicitly multivariate approaches to understanding genotype-phenotype maps are clearly needed.

This convergence of genetic effects on axes of covariation is also reflected in our finding that mutations to major developmental genes produce effects that tend to align with the directions of effect associated with the corresponding broader pathways or ontological groups. Our analysis focused on two specific processes— FGF signaling and chondrocyte differentiation.

There was a strong correlation between the *Fgf10* mutant and the FGF signaling pathway effect, while the *Bmpr1b* mutant effect was moderately correlated to the chondrocyte differentiation direction. For both process MGP maps, other mouse models in the same pathway showed significant but weaker correlations in direction of effect. These results suggest that perturbations that are developmentally similar tend to move the phenotype in the same direction in multivariate space (Figure 7B). Even so, both mutational and higher-level pathway/process effects tend to converge on a few directions of variation suggesting that multiple pathways and processes lead to common developmental outcomes. This conclusion is further supported by our finding that the genetic axes of covariance for individual processes/pathways can align with multiple directions of mutational effect. For example, the process MGP phenotypes highlighted in the white rectangle in figure 7B are all highly correlated with a set of BMP and growth hormone-related mutants.

In some cases, mutants and MGP map directions do not correspond. There are several ways this can occur. The first is that the DO population may simply lack alleles as deleterious as found in mutant lines. A small effect allele in the DO may not align with the direction of a mutant almost completely lacking expression of the target gene. Further, there are many examples where a mutation may have different and sometimes even opposite effects depending on genetic background (Mackay, 2014; Percival et al., 2017). Mutations of major effect may also differ in direction from variants in related genes that have smaller phenotypic effects due to underlying nonlinearities in development (Green et al., 2017). Investigating how variants in genes that are functionally related vary in phenotypic effect is an important avenue of inquiry that is revealed by analyses such as those we have performed here. Additionally, relationships between process and mutant effects may stimulate hypotheses about previously unknown or unvalidated interactions between loci or pathways.

A second potential reason that MGP effects may not correspond to major mutation effects is the use of only one PLS axis for each process analysis. With only one axis, we select the phenotypic direction with greatest covariance with genetic marker variation. If there are multiple large marker effects that do not covary, the weaker marker effect will be masked in the analysis. For instance, there may be a PLS axis for “chondrocyte differentiation” that corresponds more strongly with the *Bmp2* mutant phenotype (Supp fig 3). This phenomenon may be particularly prominent for pathways with substantially different mutant effects, like FGF (Fig 7A).

Finally, our analysis shares the limitation of all approaches based on gene annotation data. Incomplete annotation may contribute to lead to faulty or incomplete groupings of genes when defining pathway/process hypotheses. Gene annotation is a huge undertaking, and there is substantial variation in the completeness of different process annotations. Many process annotations are manually assigned using inference from the literature, while most are a combination of automated efforts based on transcript similarity and human curation (Mudge and Harrow, 2015). Related to this, we assign gene annotation data to genetic markers based on the closest protein-coding region. While this is a reasonable proxy, there will be regulatory sites that affect genes other than the one immediately adjacent and this is a potential source of uncertainty in our analysis.

The MGP method represents a deliberate decision to trade higher level insight from genotype-phenotype association data at the expense of statistical certainty about the significance of individual gene effects. The current implementation of the method also does not allow for quantification of individual epistatic effects. Epistasis occurs when the genotypic trait value for a locus is altered by the genotype of a different locus. Such effects generate nonlinear genotype phenotype maps, but when considered genome-wide, contribute mainly to additive variance (Cheverud and Routman, 1995; Hill, 2017). The MGP method is additive in that it models only the linear effects of genes. However, since it captures the covariances among genotypic effects, much of this “additive” variation is likely epistatic in origin.

Complex traits present a massive challenge in genomics because so many are turning out to be enormously polygenic. To generate tractable explanations of the genetic basis for such traits, methods are needed that extract higher-level representation of genotype-phenotype relationships than those that emerge from single-locus focused approaches. Here, we present an hypothesis-driven framework for deriving such higher-level genetic explanations for phenotypic variation. Our approach leverages the biological tendency for developmental processes to produce covariation among aspects of a multivariate phenotypic trait (Hallgrimsson et al., 2009; Wagner et al., 2007). The underlying assumption in this approach is that there are latent variables within high-dimensional genotype-phenotype data that correspond to developmental architecture. We believe that analyses aimed at defining and characterizing such latent variables represent a level of genetic explanation for phenotypic variation that is complementary to genetic analyses designed to establish the significance of single locus effects. Pursuing such questions will help bridge the gap between emerging mechanistic accounts of morphogenesis and our growing understanding of the genetics of morphological variation.

## Methods

### Mice

We use a sample (*n* = 1,145) of Diversity Outbred mice (DO; Jackson Laboratory, Bar Harbor, ME) to map GP relationships for craniofacial shape (Churchill et al., 2012, 2004). The DO is a multiparental outcross population derived from the eight founding lines of the Collaborative Cross (CC). Each animal’s genome is a unique mosaic of the genetic diversity found in the CC— more than 45 million segregating SNPs (Consortium, 2012). Random outcrossing over many DO generations maintains this diversity and, with recombination, increases mapping resolution.

Our DO sample was sourced from three separate laboratories and seven DO generations. 386 are from the Jackson Laboratory (JAX), 287 from the University of North Carolina (UNC), and 472 come from the Scripps Research Institute. Supplemental figure 4 shows the distribution of the sample by lab source and generation of breeding. Imaging of mice at the University of Calgary was performed under IACUC protocol AC13-0268. Ankrd11 and Bmpr1b mutant mice were bred at the University of Alberta by the Graf lab under Animal Use and Care Committee protocol AUP1149, in accordance with guidelines of the Canadian Council of Animal Care.

### Genotyping

Genotyping was performed by Neogen (Lincoln, NE). Ear clippings were used to extract DNA for all samples. Mice from generations 9, 10, and 15 were genotyped using the MegaMUGA genotyping array (77,808 markers); mice from generations 19, 21, 23, and 27 were genotyped using the larger GigaMUGA array (143,259 markers) (Morgan et al., 2016). To pool the genotype data from these two SNP arrays with differing numbers of markers, we imputed markers between the two genotyping arrays using the “calc_genoprob” function in the qtl2 package (Broman et al., 2018). The function uses a hidden Markov model to estimate genotype probabilities and missing genotype data (Gatti et al., 2014). After imputation, the merged genetic dataset consists of 123,309 SNPs which vary among CC founders. Each animal’s genetic record is a 123,309*8 matrix of estimated diplotype contributions of each CC founder to each marker.

### Scanning and landmarking

We used micro-computed tomography to acquire 3D scans of the full heads of the mice. Scanning was done at the University of Calgary at .035 mm voxel resolution (Scanco vivaCT40). One of us (WL) then acquired 54 3D landmarks (Fig 9) manually on each volume using Analyze 3D. A discussion of the error associated with manual landmarking can be found in Percival et al (Percival et al., 2019).

**Fig 9.**
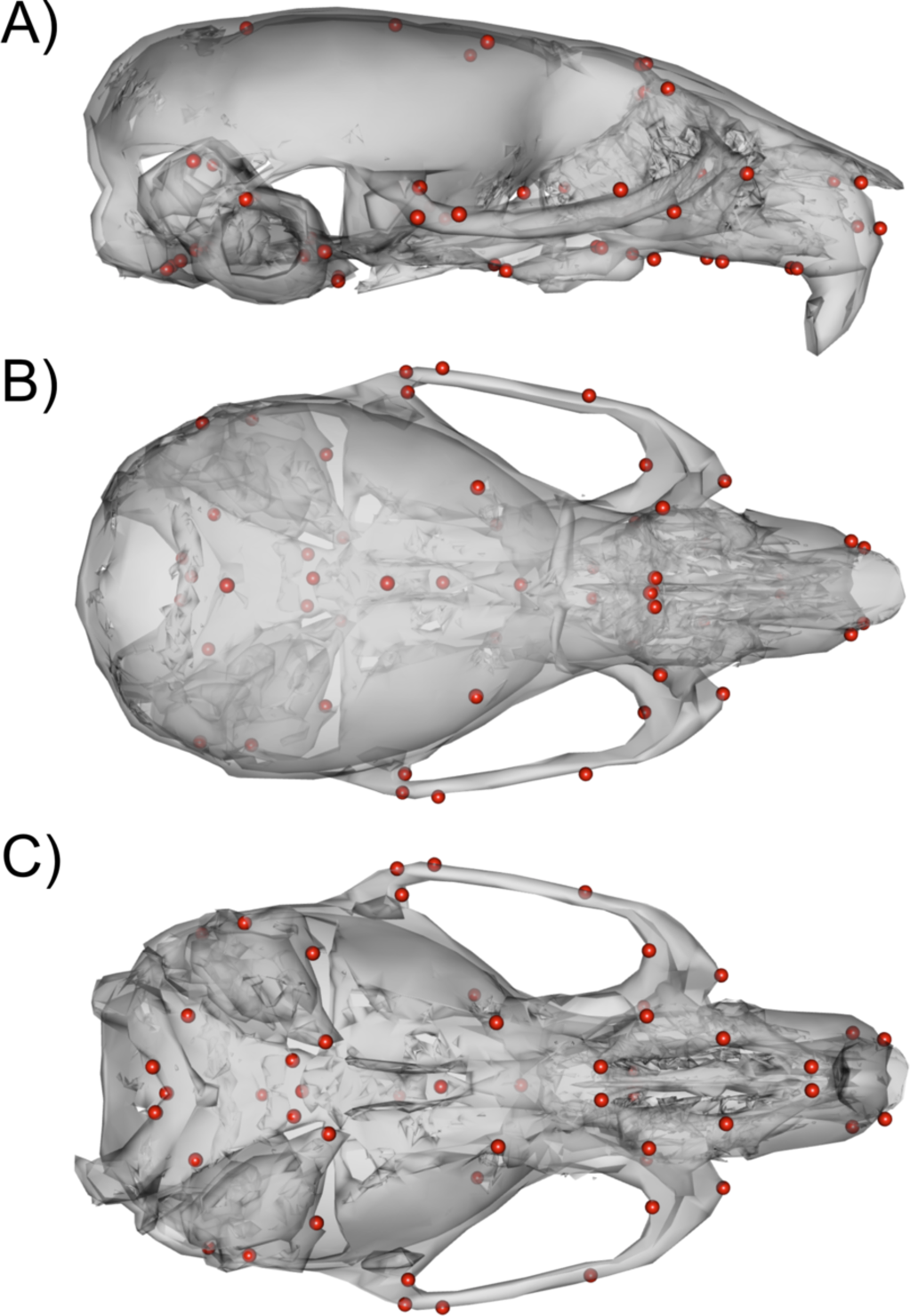
54 3D landmark configuration. A) Sagittal view of representative scan with landmarks shown as red spheres. B) Dorsal view of landmark configuration. C) Ventral view of landmark configuration.

### Landmark registration

We symmetrized landmarks along the midline of the skull using Klingenberg et al.’s method for object symmetry which configures landmark pairs into a common orientation with reflection and subsequently removes variation associated with translation, scale, and rotation, using Generalized Procrustes Analysis (Adams and Otárola-Castillo, 2013; Klingenberg et al., 2002; Mardia, 2000; Schlager, 2017). To focus on shared, within-generation patterns in our multigenerational DO sample, we regressed symmetric shape on DO generation, and used the residual shapes with the grand mean added as the observations for analysis.

### Genetic relatedness

Adjustment of phenotypes for the influence of genetic relatedness is a common approach in genomic studies to prevent spurious associations. However, it is not necessary in all cases, such as situations with low genetic relatedness and little variation in relatedness. We evaluated whether accounting for genetic relatedness was important for our sample. To do so, we estimated a kinship matrix based on DO genotype correlations (Cheng et al., 2013; Broman et al., 2019).

The kinship values in our sample have a mean of 0 and a standard deviation of .047 (Supp fig 5). As a result of these findings, we performed all subsequent analyses on the within-generation symmetric shape data, without an adjustment for relatedness.

### Regularized PLS analysis

Multivariate genotype-phenotype methods for explicitly modeling multivariate phenotypes and for overcoming the limitations of simple linear regression are increasingly common in mapping studies. Claes et al. (Claes et al., 2018) used canonical correlation analysis to quantify individual SNP effects for a multivariate measurement of facial shape. Each test returns a vector of the linear combination of phenotypic effects that maximally correlates to the alleles at a given locus. Mitteroecker et al. (Mitteroecker et al., 2016) developed a multivariate strategy around a singular value decomposition (SVD) of GP covariation. Partial least squares (PLS) describes a family of approaches that use SVD to decompose cross covariance matrices (Lee et al., 2011; Mitteroecker and Gunz, 2009; Singh et al., 2016). PLS is increasingly used with large genetic datasets in order to model how genomic effects extend to multiple traits (BJØRNSTAD et al., 2004; Mehmood et al., 2011; Tyler et al., 2017). However, its implementation for MGP mapping is, thus far, much more limited.

SVD decomposes the covariance matrix into three matrices:

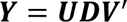

Where **Y** is the mean-centered covariance matrix, **U** denotes the left singular vectors, a set of vectors of unit length describing the relative weighting of each variable on each axis, and **D** denotes the variance along each axis. **V** denotes the set of right singular vectors. For a full (square, symmetric) covariance matrix, **U** and **V** are identical, and the decomposition is equivalent to PCA. For a non-symmetric matrix of covariances, i.e., one describing covariance between two distinct blocks of traits, each successive column of **U** and **V** provide a pair of singular vectors describing the best least squares approximation of covariance between the two blocks, in order of greatest covariance explained to least.

PLS is most often used to find low-rank linear combinations that maximize covariance between two sets of features. Here, we use a data-driven regularized PLS model implemented in the mddsPLS package to find paired axes that maximize covariance between allelic and shape variation (Lorenzo et al., 2019). The model uses a lasso penalty to minimize the coefficients (loadings) towards zero to prevent overfitting (James et al., 2013). Overfitting can occur in when many genotypic markers are included in the model, particularly when markers are colinear. The genotype block is composed of the full set of DO founder probabilities for each selected marker. Thus, an analysis of 20 markers would estimate 160 genotype coefficients. The phenotype block consists of the full set of 54 3-dimensional landmarks (162 phenotype coefficients). In all biological process analyses undertaken herein, we used a regularization parameter of 0.06 and report only the first paired axes of the PLS model, i.e., the genotype and phenotype axes which explain the most covariance.

### Biological process gene sets

For process-specific MGP analyses, we used the mouse genome informatics database (Bult et al., 2018) to identify genes annotated to a given process. Each annotation term has an associated GO ID. For example, “chondrocyte differentiation” has GO ID GO:000206 (Fig 10, box 1). We cross-reference the GO ID with the Ensemble genome database (GRCm38.p6) to find the name, chromosome, and base pair start/end position for each gene (Fig 10, box 2) annotated to the process. For genes with multiple splice variants, we select the full transcript start/end positions. For each gene, we compare marker base pair positions and select the closest upstream and downstream markers to the center of each gene. The 8-state genotype probability is then calculated as the average founder allele probabilities between the two selected markers. (Fig 10, box 3). After marker selection, we fit the regularized PLS model using the founder allele probabilities (8 variables/marker) and full landmark data set (Fig 10, box 4).

**Fig 10.**
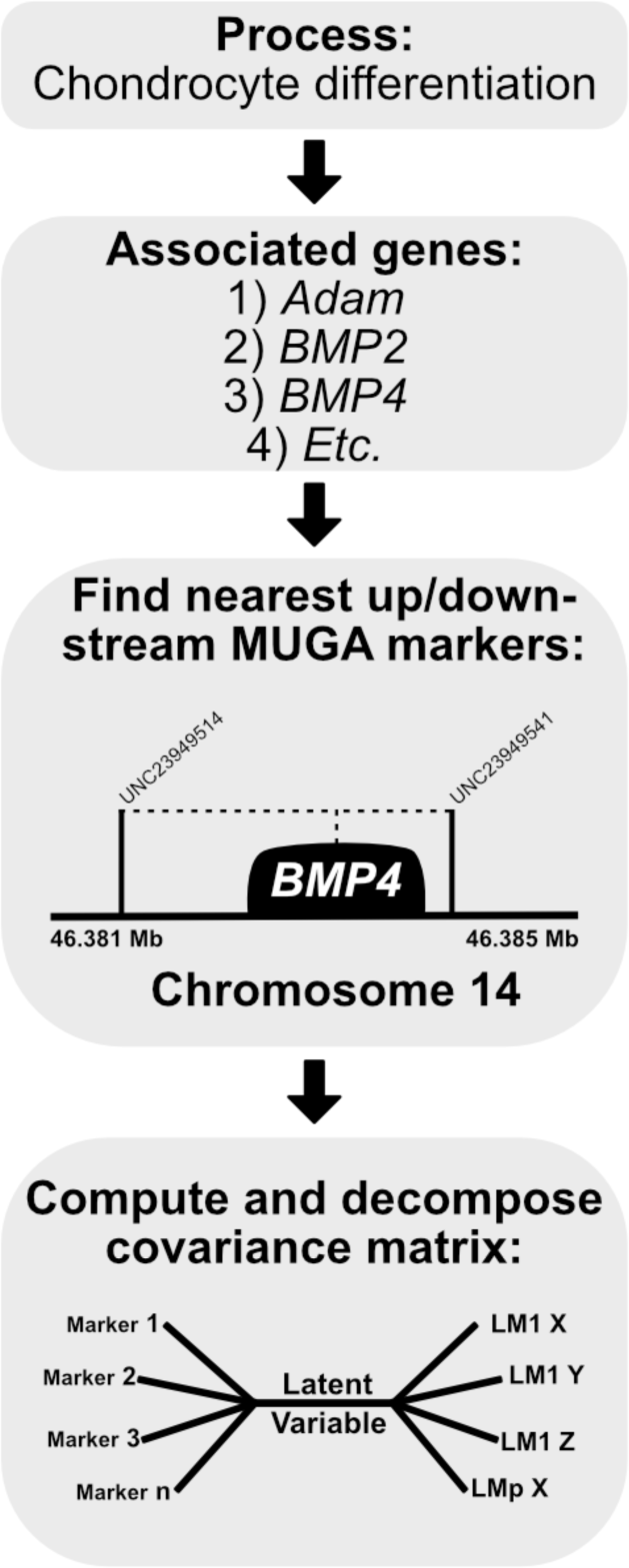
Process MGP schematic. Once a process is selected, we cross-reference the known gene locations with the locations of the genotyped markers in the DO sample. The founder probabilities of the nearest upstream and downstream markers are averaged for each gene. The compiled founder probabilities and landmark coordinates are then used in a regularized PLS model to estimate latent axes of covariation.

We generate graphical displays of process results using the R packages ggplot2 (Wickham, 2016) and Morpho (Schlager, 2017). An example script to reproduce the analyses is provided at **github.com/j0vid**.

### Statistical results and comparisons

We estimate the magnitude and direction of MGP process effects using R^2^ and vector correlations, respectively. R^2^ is calculated as the ratio of trace of the predicted model covariance to the trace of the phenotypic covariance matrix. We contextualize the MGP process R^2^ by comparing it to the R^2^ value of 1000 randomly drawn marker sets of the same size. For instance, a process annotated with 40 genes would be compared to 1000 40-gene MGP analyses with random markers selected in each iteration. Random marker selection for permutation is constrained to follow similar patterns of linkage disequilibrium to the observed marker set of interest. The null expectation in this scenario is that gene annotation does not provide better information about coordinated marker effects than a randomly selected set of markers.

Vector correlations between process MGP effects are calculated by taking the Pearson product-moment correlation of the two sets of process PLS1 phenotypic loadings. Vector correlations between process effects and mutant effects are calculated by taking the correlation between the process PLS1 phenotypic loadings and mutant MANOVA coefficients. The MANOVA compares the mutant group phenotype with the DO sample specified as the reference group.

### Chondrocyte morphometrics

Chondrocyte morphometrics were performed using a novel technique developed by the Marcucio laboratory. Images of the intersphenoid synchondrosis (ISS) were stained with H&E, SafO, or picrosirius red were captured and imported into ImageJ (2-6 sections from at least 4 mice/genotype/synchondrosis). Landmarks were placed in a defined order (left, right, top, bottom) of visible chondrocytes in the synchondrosis using the ImageJ’s multi-tool. Data points were then exported as XY coordinates and imported into Microsoft Excel for calculation of major and minor axes relative to overall width of synchondrosis. Area of individual cells was determined from height and width values based on assumption that each cell is roughly ellipsoidal. An example of major and minor axis measurements and ellipsoidal area measurements on a slide is provided in supplemental figure 6.

We compared differences in the distribution of cell sizes along normalized synchondroses between *Bmpr1b* mutants and controls with a mixed effects model approach. We used ellipsoidal area of cell size (in microns) as our dependent variable. For fixed effects, we modelled the normalized synchondrosis position (1^st^ and 2^nd^ order), where a value of 0 represents the relative midline of the synchondrosis and values of -1 and 1 represent the most distant cells in that synchondrosis. We also modelled genotype as a fixed effect as well as a genotype by cell position interaction (both 1^st^ and 2^nd^ order interactions). For each individual within each genotype, we measured multiple histological sections. These repeated and nested measurements of cell size in multiple sections for each individual were modelled as random effects. To test for cell size differences between genotypes, we used a likelihood ratio test to compare the full model to a reduced model with the fixed effect of genotype removed.

### Visualization tools

We introduce an interactive web application that allows the user to select processes of interest with a graphical user interface and see the resulting craniofacial effect at **genopheno.ucalgary.ca/MGP**. The web apps were written using the shiny package in R (Chang et al., 2018). The application dynamically filters the MGI GO database based on the initial user input. Queries will only list GO terms with exact matches. For example, “chond” will return GO terms that incorporate either “chondrocyte” and “mitochondria”.

Multiple queries can be selected. An analysis of “chondrocyte differentiation” and “chondrocyte hypertrophy” will select the joint gene set of both processes. Processes with different names can be jointly queried with the pipe operator “|”, which is interpreted as an OR (union) operator. For example, to generate the list of GO terms associated with either apoptosis or WNT, we used the “apoptosis|WNT” query and selected the processes “Wnt signaling pathway” and “execution phase of apoptosis” to perform the analysis on the joint set of associated genes (Supp fig 7).

Several other parameters can be specified by the user including the type of plot to be generated for the genetic loadings, the amount of magnification applied to the phenotype effect vectors, the regularization parameter, and the option to overlay a mutant phenotype for comparison. The comparative database currently includes craniofacial shape contrast data (wild-type vs. mutant) for 30 mutant genotypes. If a mutant comparison is selected, the full set of DO specimens are registered with the mutants added (with size removed). We then provide the vector correlation between the process effect and the mutant effect (see Fig 8). The database also includes PC1 of the DO sample for comparison.

The app enables users to save results. A save request will generate and download an HTML report of the analysis which includes several versions of the genetic effect plot and an interactive 3D model of the estimated phenotypic effect. If a mutant comparison is selected, it will also appear in the report.

The application tracks recent searches by the user for their reference. A heatmap of process vector correlations of the PLS phenotype loadings is also available under the “recent searches” tab. The user can select between a heatmap of the processes in their search history or a random assortment of process correlations from past, anonymous user searches.

## Acknowledgements

Grants: NIH - 2R01DE019638 to RM, BH and JC, NSERC 238992-17, CIHR Foundation grant 159920 to BH, CFI grant #36262 to BH and NSERC RGPIN-2014-06311 to DG. JDA is supported by an Eyes High fellowship, an Alberta Children’s Hospital Research Institute scholarship and a MITACS graduate fellowship.

## Supplemental Material

**Supplemental figure 1.**
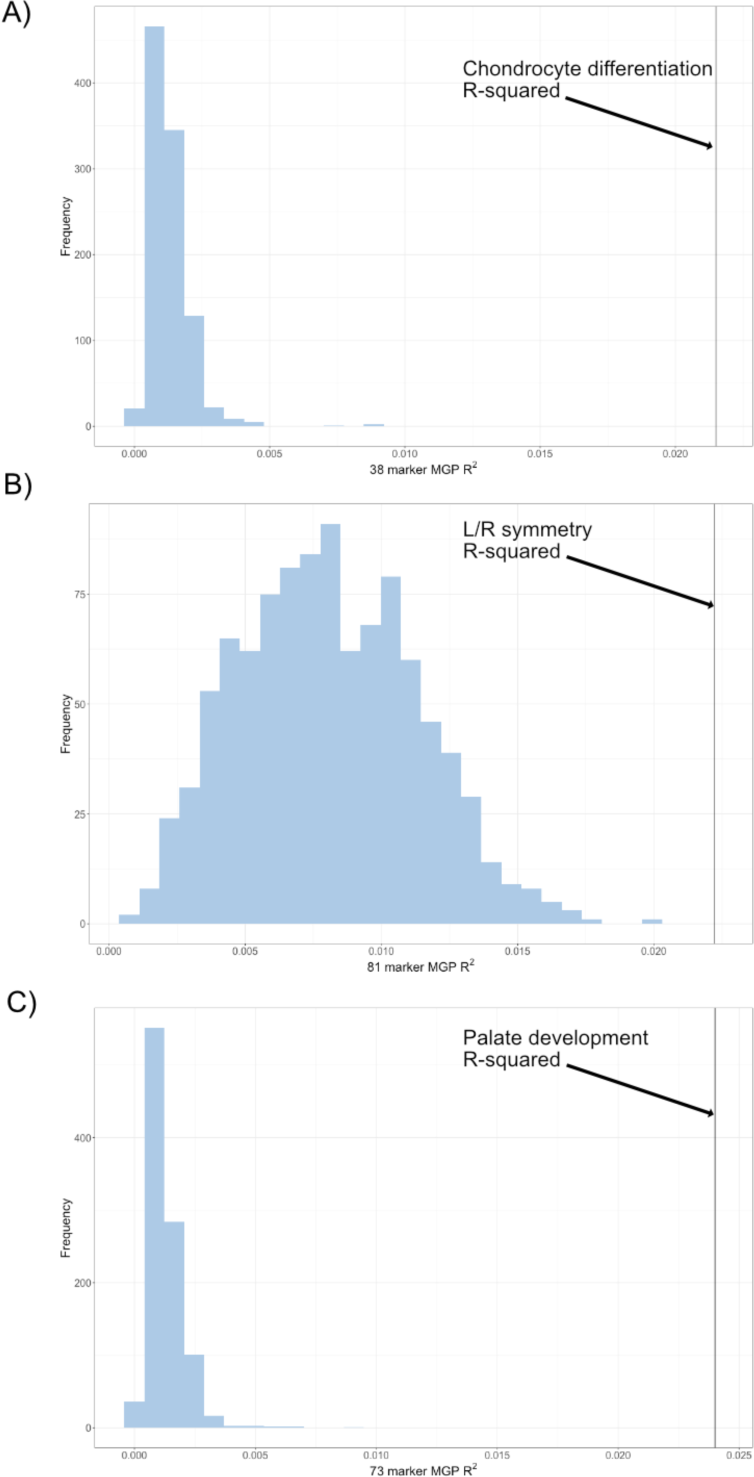
Permutation of marker sets of fixed size. A) The permuted R^2^ distribution of 1000 38-marker MGP analyses is shown in blue. The estimated R^2^ of the chondrocyte differentiation MGP is shown as a black vertical line. B) The permuted R^2^ distribution of 1000 81-marker MGP analyses is shown in blue. The estimated R^2^ of the L/R symmetry MGP is shown as a black vertical line. C) The permuted R^2^ distribution of 1000 73-marker MGP analyses is shown in blue. The estimated R^2^ of the palate development MGP is shown as a black vertical line.

**Supplemental figure 2.**
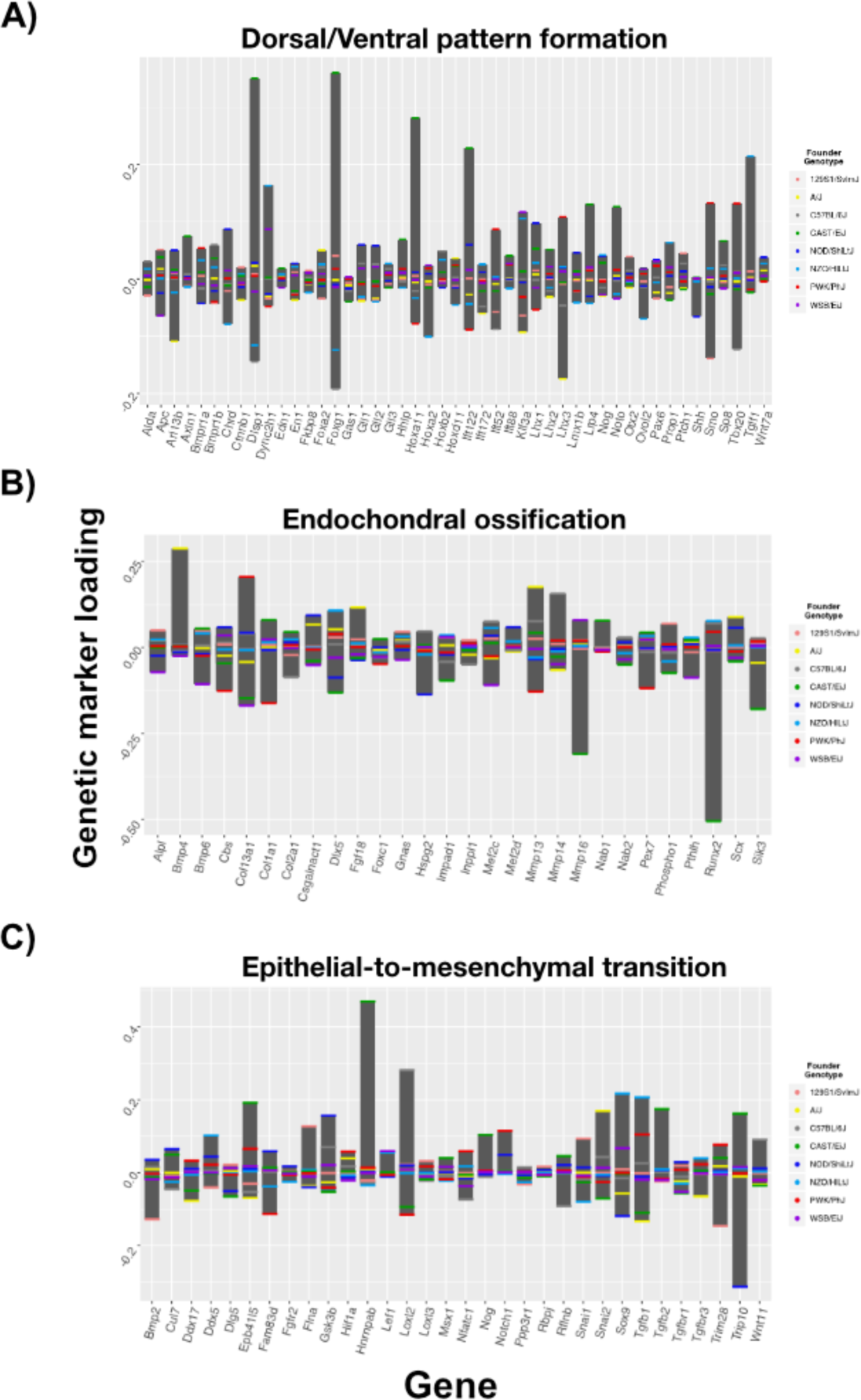
Genetic effect loadings for multiple process MGPs. Selected processes are listed above their respective plots. Marker loadings are not scaled to a common range.

**Supplemental figure 3.**
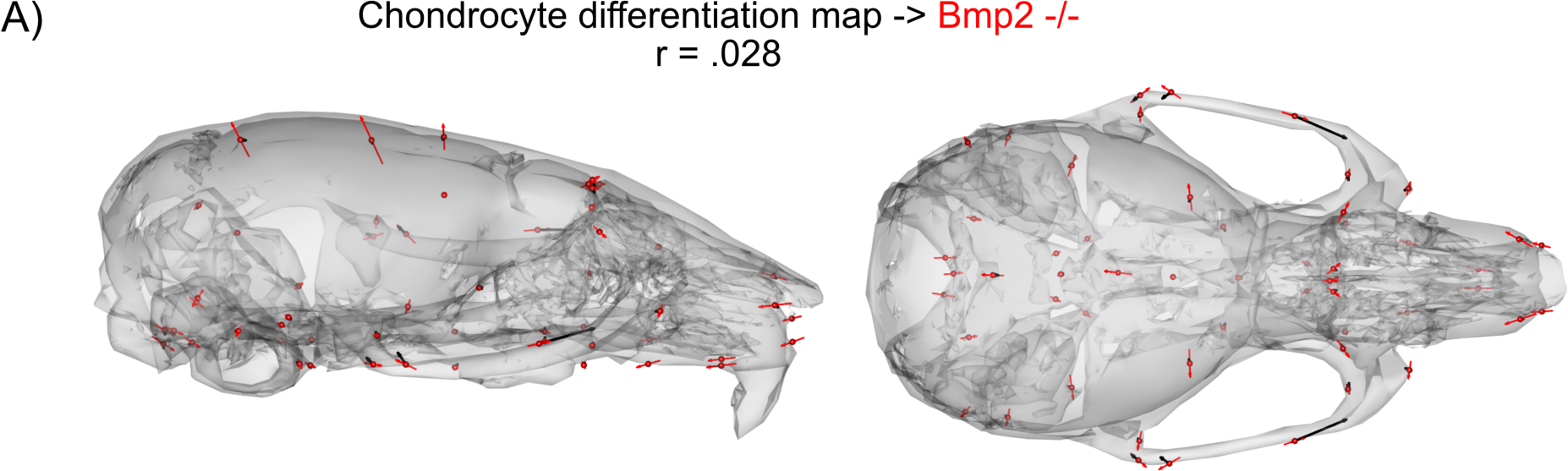
Additional process-to-mutant comparisons. A) The estimated chondrocyte differentiation MGP phenotype magnified 4x (black) with *Bmp2* homozygous mutant (red).

**Supplemental figure 4.**
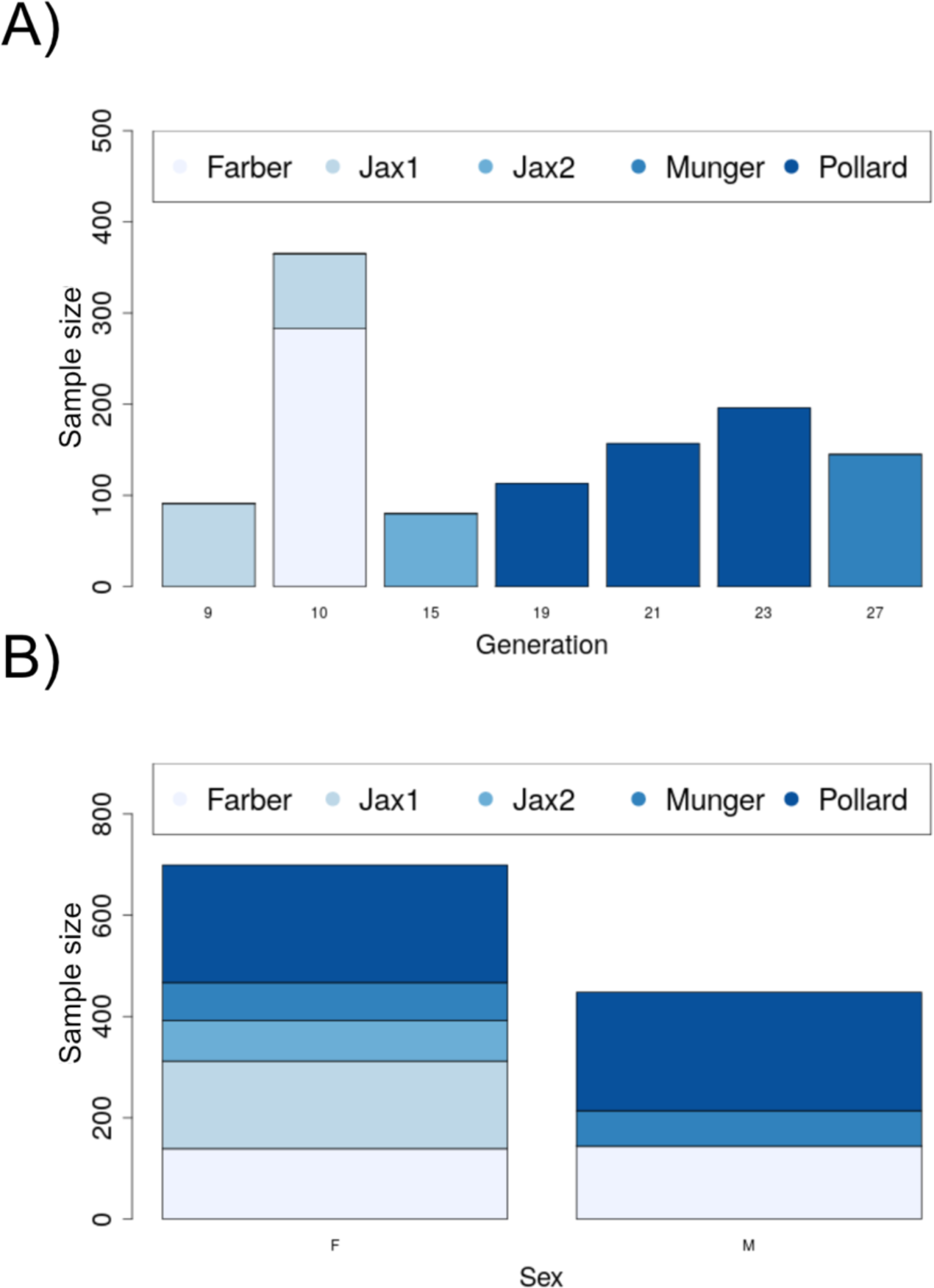
Demographic plots for the DO sample. A) The distribution of the sample by generation and data source (lab). B) Distribution of sex by source (lab).

**Supplemental figure 5.**
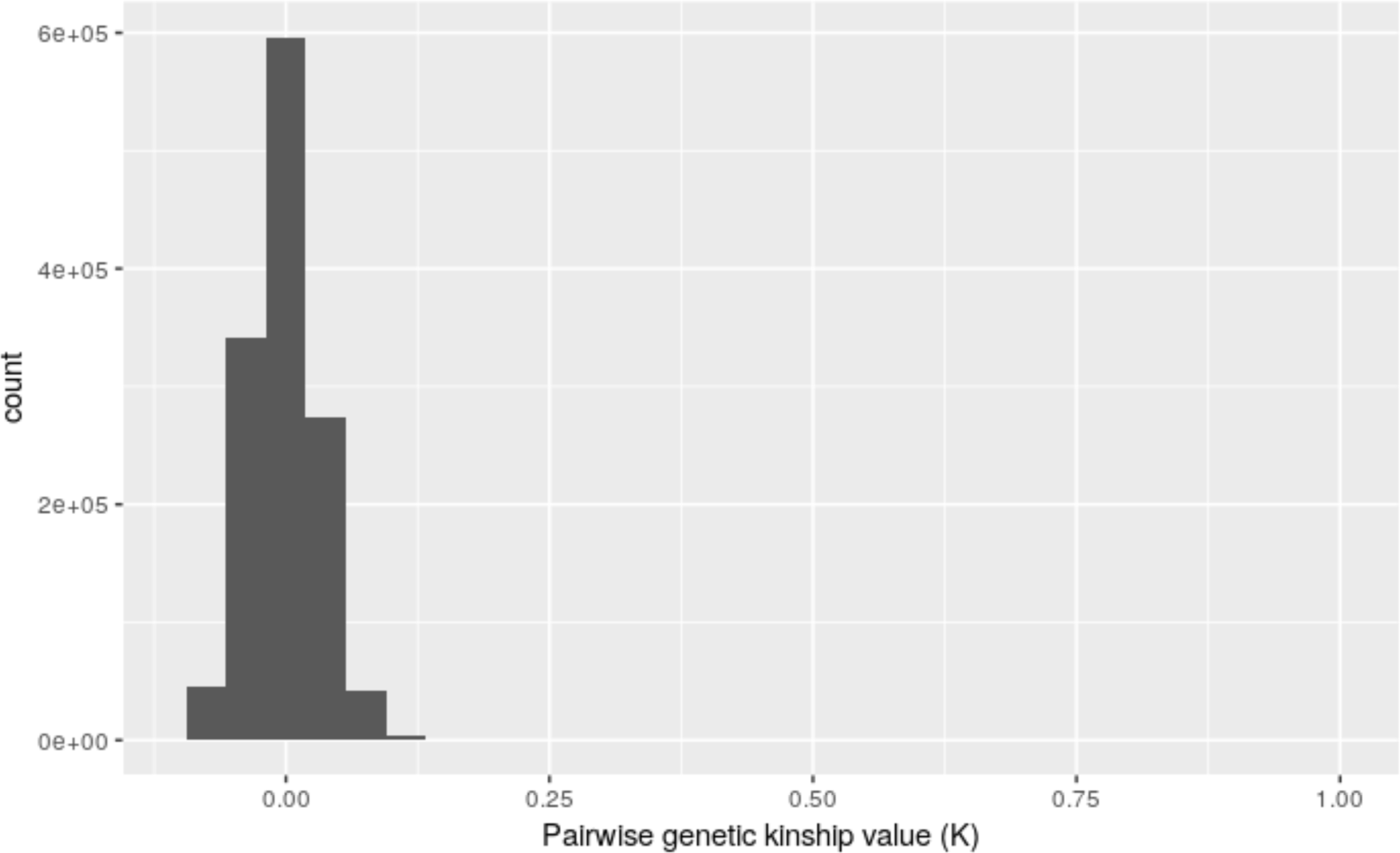
Histogram of kinship matrix values in the DO sample.

**Supplemental figure 6.**
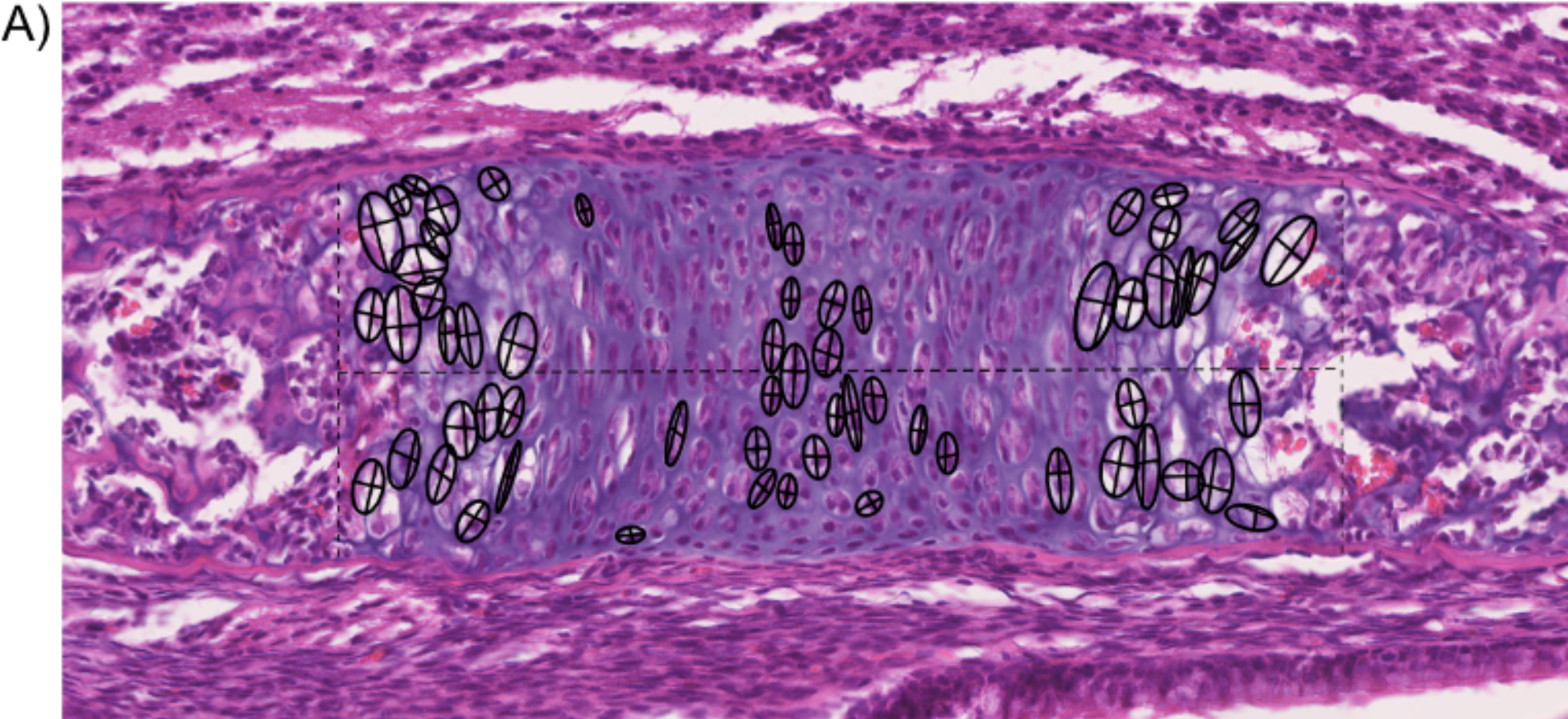
Chondrocyte morphometric example. Landmarks are placed in the top, bottom, left, and right sides of the cell to best capture the height and width of the cells (show here as crosses). The height and width measurements are then used to calculate the area of an ellipsoid as an approximation of cell size.

**Supplemental figure 7.**
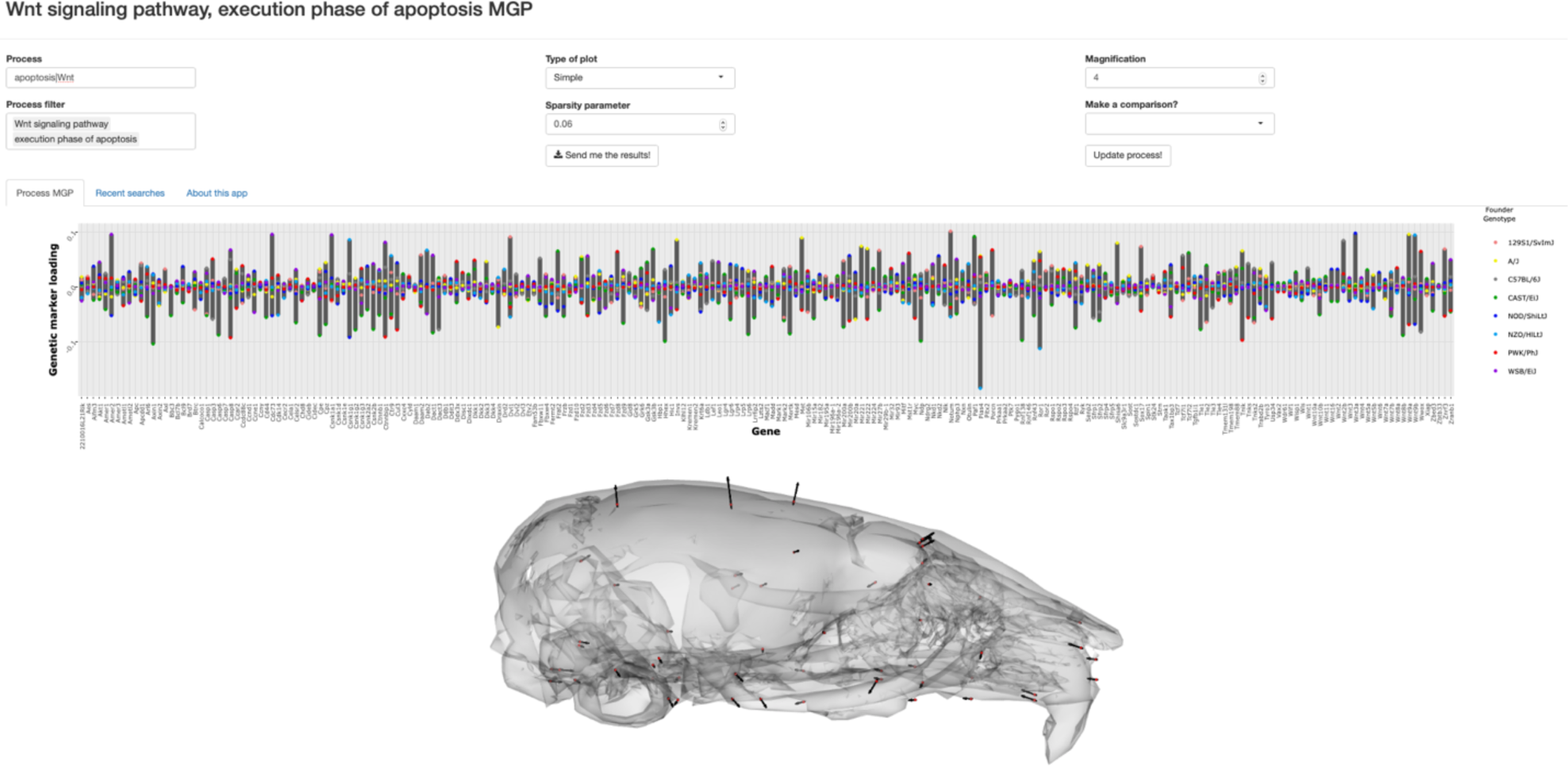
Combining queries in the MGP shiny app with the pipe operator. In order to filter the GO database with multiple terms, the pipe operator can be used as shown. Here, the user has selected processes associated with either the apoptosis or Wnt pathway process. The barplot shows the relative effect sizes for markers associated to both “Wnt signaling pathway” and “execution phase of apoptosis” GO terms.

**Supplemental table 1.**
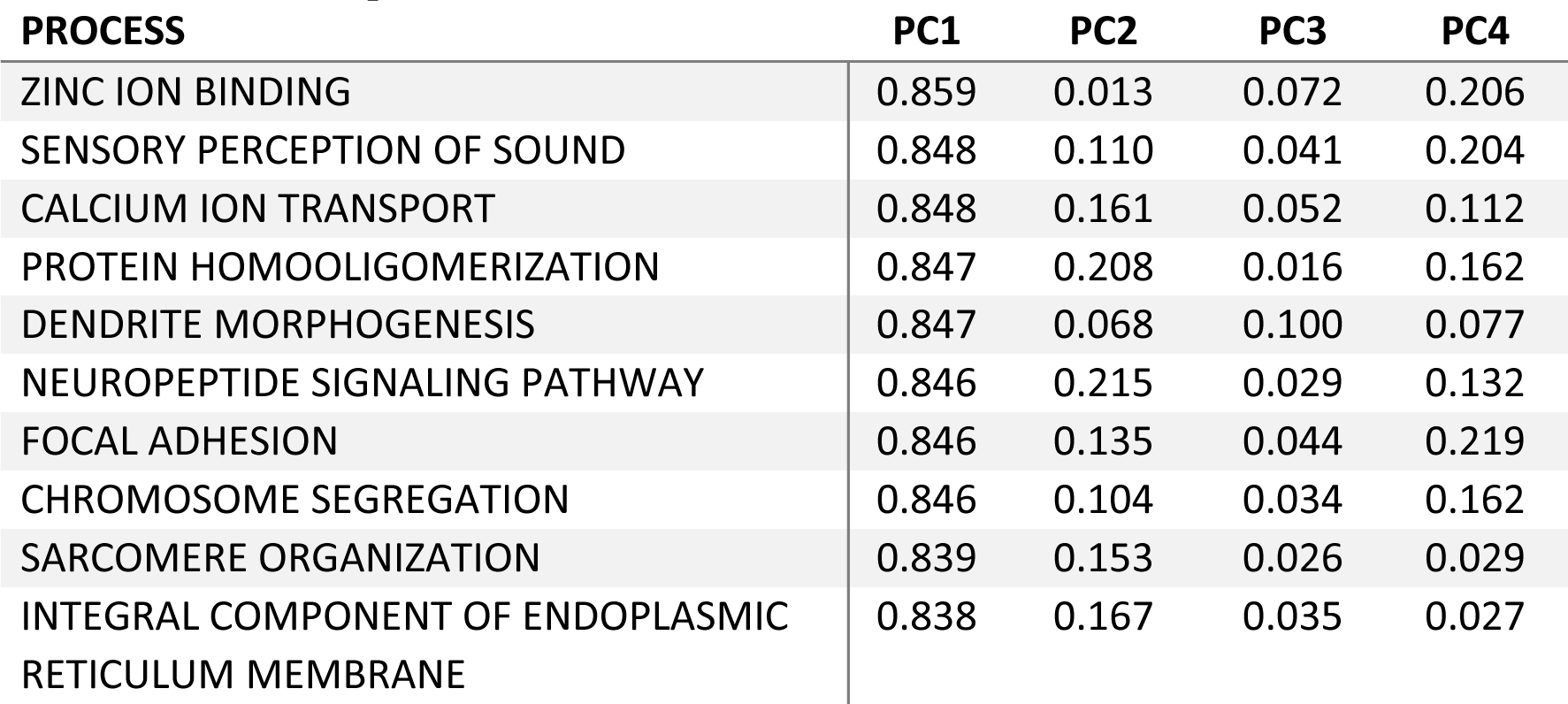
Top 10 MGP vector correlations with PC1. Corresponding PC2-4 vector correlations are also provided.

**Supplemental table 2.** Processes with moderate vector correlations to PC 1-4.

| **PROCESS** | **PC1** | **PC2** | **PC3** | **PC4** |
| --- | --- | --- | --- | --- |
| NEGATIVE REGULATION OF I-KAPPAB KINASE/NF-KAPPAB SIGNALING | 0.541 | 0.256 | 0.492 | 0.259 |
| LOCOMOTORY EXPLORATION BEHAVIOR | 0.380 | 0.211 | 0.689 | 0.241 |
| AU-RICH ELEMENT BINDING | 0.570 | 0.330 | 0.403 | 0.208 |
| TRANSCRIPTION REGULATORY REGION SEQUENCE-SPECIFIC DNA BINDING | 0.701 | 0.093 | 0.444 | 0.263 |
| DYNEIN COMPLEX | 0.685 | 0.246 | 0.314 | 0.194 |
| PRESYNAPTIC CYTOSOL | 0.392 | 0.383 | 0.135 | 0.508 |
| NEGATIVE REGULATION OF FAT CELL DIFFERENTIATION | 0.391 | 0.245 | 0.730 | 0.030 |
| CELL BODY | 0.802 | 0.251 | 0.069 | 0.263 |
| ADULT BEHAVIOR | 0.746 | 0.392 | 0.035 | 0.210 |
| PROTEASOMAL UBIQUITIN-INDEPENDENT PROTEIN CATABOLIC PROCESS | 0.617 | 0.400 | 0.010 | 0.263 |

